# Microbial community and geochemical analyses of trans-trench sediments for understanding the roles of hadal environments

**DOI:** 10.1101/729517

**Authors:** Satoshi Hiraoka, Miho Hirai, Yohei Matsui, Akiko Makabe, Hiroaki Minegishi, Miwako Tsuda, Juliarni, Eugenio Rastelli, Roberto Danovaro, Cinzia Corinaldesi, Tomo Kitahashi, Eiji Tasumi, Manabu Nishizawa, Ken Takai, Hidetaka Nomaki, Takuro Nunoura

## Abstract

Hadal trench bottom (>6,000 m below sea level) sediments harbor higher microbial cell abundance compared to adjacent abyssal plain sediments. This is supported by the accumulation of sedimentary organic matter (OM), facilitated by trench topography. However, the distribution of benthic microbes in different trench systems has not been explored yet. Here, we carried out small subunit ribosomal RNA gene tag sequencing for 92 sediment subsamples of seven abyssal and seven hadal sediment cores collected from three trench regions in the northwest Pacific Ocean: the Japan, Izu-Ogasawara, and Mariana Trenches. Tag-sequencing analyses showed specific distribution patterns of several phyla associated with oxygen and nitrate. The community structure was distinct between abyssal and hadal sediments, following geographic locations and factors represented by sediment depth. Co-occurrence network revealed six potential prokaryotic consortiums that covaried across regions. Our results further support that the endogenous OM cycle is driven by hadal currents and/or rapid burial shapes microbial community structures at trench bottom sites, in addition to vertical deposition from the surface ocean. Our trans-trench analysis highlights intra- and inter-trench distributions of microbial assemblages and geochemistry in surface seafloor sediments, providing novel insights into ultra-deep-sea microbial ecology, one of the last frontiers on our planet.

## Introduction

The abyssal plain extends from the continental slope to the rim of deep trenches (3,000–6,000 m below sea level [mbsl]) and covers 85% of the global seafloor area, while the hadal zone (>6,000 mbsl) comprises 1–2% of it [1, 2]. In general, abyssal water and sediments are usually oligotrophic, and physical and chemical conditions (*e.g.*, salinity, temperature, dissolved oxygen, and nutrient concentrations) in hadal water are similar to the overlying abyssal water despite the higher hydrostatic pressure [1–3]. However, cell abundance and microbial carbon turnover rates are significantly higher at hadal compared to abyssal seafloors, especially below the sediment surface, while microbial abundance in surface sediments are sometimes comparable between hadal and abyssal sites [4]. This could be attributed to factors apart from differences in the vertical downward flux of sinking organic matter (OM) from the ocean surface and hydrostatic pressure.

Hadal zones are generally located in oceanic trenches that are formed along plate boundaries by movement of oceanic plates, and thus experience episodic and/or regular landslides [5, 6]. These landslides cause downward transportation of surface sediments along with relatively fresh OM via the funnel effect of trench geomorphology [7–11]. Moreover, higher sedimentation rates and concentrations of subseafloor organic compounds in trench bottom sediments compared to neighboring abyssal sediments have been reported in multiple trench regions under oligotrophic and eutrophic oceans [4, 11–14]. Therefore, the intrinsic organic carbon sources in hadal zones are considered to facilitate establishment of distinctive faunal and prokaryotic biospheres observed at global scales [15–19]. Recently, the influence of physicochemical features on hadal biospheres were reported for microbial communities in the Mariana and the Kermadec Trench regions under oligotrophic oceans [20, 21]. However, except for the cases, microbial community structures in hadal sediments were not compared with adjacent abyssal regions [2], and such comparison under eutrophic oceanic regions have not been investigated yet.

Here, we evaluated prokaryotic community structure in 92 sediment subsamples of 14 sediment cores collected from four hadal and seven adjacent abyssal sites located in three trenches in the northwest Pacific Ocean; the Japan, Izu-Ogasawara (Izu-Bonin), and Mariana Trenches (Fig. 1 and Table S1). The Japan and Mariana Trenches lie under eutrophic and oligotrophic oceans, respectively, while the primary productivity of the ocean above the Izu-Ogasawara Trench presents intermediate features. We performed geochemical analyses of the sediments and culture-independent microbial analyses including direct cell count, quantitative polymerase chain reaction (qPCR), and tag-sequencing for small subunit ribosomal RNA (SSU rRNA) gene, to investigate intra- and inter-trench diversities of prokaryotic communities and potential metabolic interactions using co-occurrence networks. Our analyses illuminated the ubiquitous existence of distinctive prokaryotic assemblages in abyssal and hadal subsurface sediments, providing new insights into the understudied oceanic microbial ecology.

**Fig. 1.**
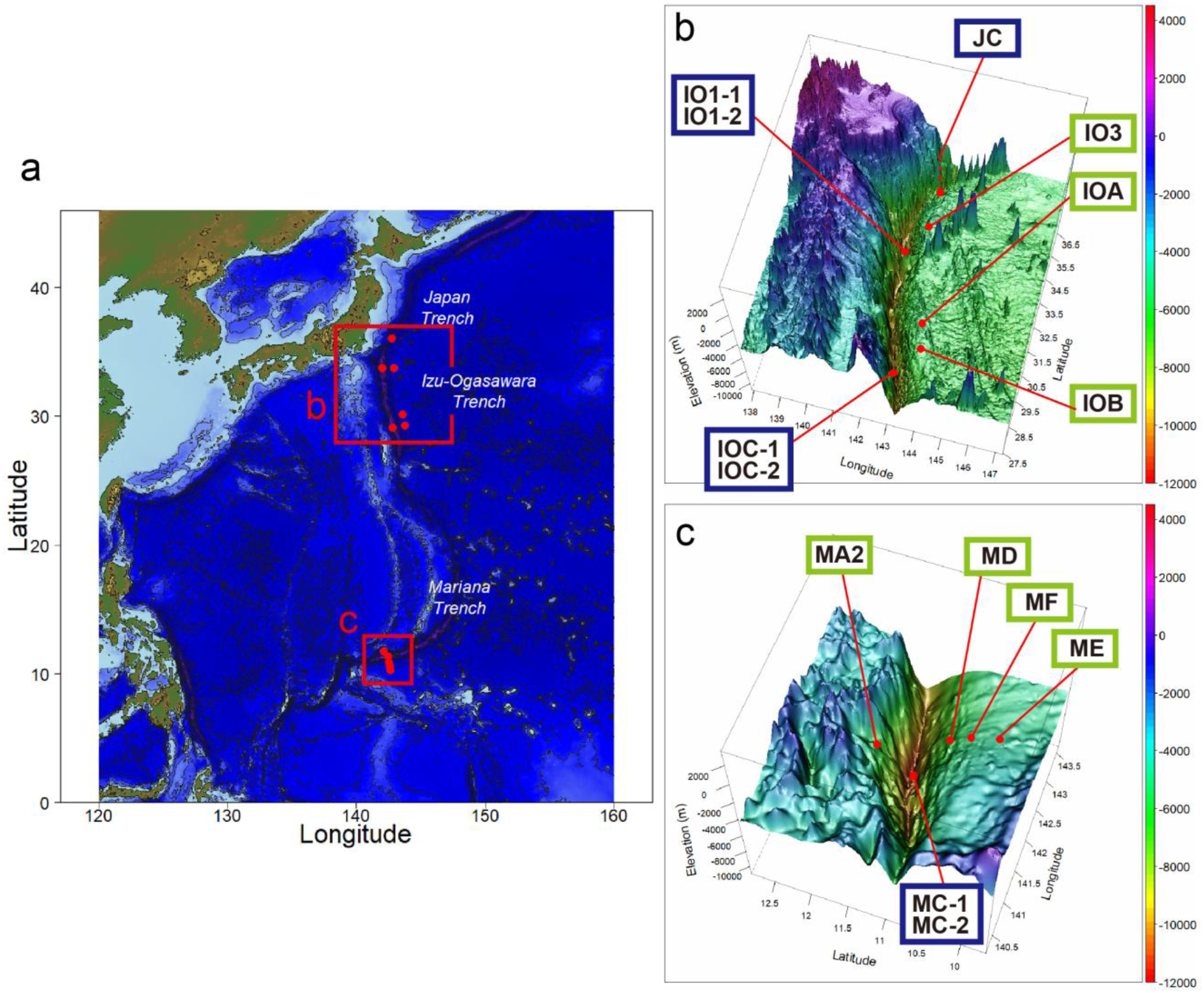
Plane (**a**) and three-dimensional (**b**,**c**) maps of the sampling stations of sediment cores and their geographic locations. Stations with green and blue rectangles represent abyssal and hadal sites, respectively. The dashed line of JC station indicates that the station is located behind the seamount.

## Materials and methods

### Sediment samplings

Sediment cores were collected over five cruises by Japan Agency for Marine-Earth Science and Technology (JAMSTEC); Research Vessel (R/V) *Kairei* KR11-11 (Dec. 2011) and KR15-01 (Mar. 2015) cruises and R/V *Yokosuka* YK11-06 (Sep. 2011) cruise for the Izu-Ogasawara Trench, R/V *Kairei* KR12-19 (Dec. 2012) cruise for the Japan Trench, and R/V *Kairei* KR14-01 cruise (Jan. 2014) for the Mariana Trench (Fig. 1 and Table S1). The water depths ranged from 4,700 to 10,902 mbsl. Among the 11 sites, 4 were located on the trench bottom (‘hadal sites’), while the other 7 were located on the trench edge (‘abyssal sites’).

Sediment cores were obtained by a gravity corer of the remotely operated vehicle (ROV) ‘*ABISMO*’ [22] during KR11-11 and KR15-01 cruises, a free fall 11K lander system [23] during KR11-11, KR12-19, and KR15-01 cruises, a multiple corer system during KR14-01 cruise, and push corers of human occupied vehicle (HOV) ‘*Shinkai 6500*’ during YK11-06 cruise. In each sampling operation, a Conductivity, Temperature, and Depth (CTD) sensor SBE-19 or SBE-49 (Sea-Bird Electronics, Bellevue, WA, USA) was used. Among the sampling operations, two operations were conducted for sites IO1 and IOC in the Izu-Ogasawara Trench (station pairs IO1-1 and IO1-2, and IOC-1 and IOC-2, respectively) and site MC in the Mariana Trench (MC-1 and MC-2). Sediment core lengths taken by the push corer, multiple corer, and 11K lander system ranged from 25 to 50 cm, while those collected from IOB and IOC-1 stations using the gravity corer were approximately 120 to 150 cm. Note that sediment-water interface of the sediment cores taken by the gravity corer might be more disturbed by sampling operations compared to the other coring systems.

Collected sediment cores were immediately subsampled onboard at 2- to 10-cm depth intervals for geochemical and microbiology analyses. Porewater was extracted by centrifugation at 2,600 × g for 10 min, then supernatant (seawater) was filtered using a 0.2-µm syringe cartridge filter. Subsamples were stored at −80 and −20°C for molecular and geochemical analyses, respectively. Sediment samples for direct cell counting were fixed with or without 5 mL formaldehyde solution (final concentration 2% w/v in PBS buffer) for 1 h at room temperature onboard and stored at −80 °C.

### Geochemical analyses

Dissolved oxygen (O_2_) concentrations in the sediment cores were measured onboard immediately after core recovery using a planar optode oxygen sensor Fibox 3 (PreSens, Regensburg, Germany). For the gravity core samples, small (∼3 mm) holes were opened along the side of the core tube at 2–5 cm depth intervals, and the sensor was inserted into the sediments. For sediment cores collected by the push corer, multiple corer, and 11K lander system, the sensor spots were attached to the inside of the transparent polycarbonate core liner tube and oxygen concentrations were measured from the outside. The sensors were calibrated with air-saturated and oxygen-free seawater before measurements. Porewater nutrient concentrations (NO_3_ ^−^, NO_2_ ^−^, PO_4_, and NH_4_ ^+^) were analyzed spectrophotometrically using an automated continuous–flow QuAAtro 2-HR analyzer (BL TEC, Osaka, Japan) in an onshore laboratory [24].

Total organic carbon (TOC), total nitrogen (TN), and their C and N isotopic compositions were measured by flash combustion with an elemental analyzer Flash EA 1112 series coupled via ConfloIV to an isotope ratio mass spectrometer DELTAplus Advantage (Thermo Fisher Scientific, Waltham, MA, USA) for KR15-01 sediment samples or DELTA V Advantage (Thermo Fisher Scientific) for the other sediment samples. Before measuring, the sediment samples were freeze-dried and acidified with 1 M HCl to remove inorganic carbon. The decalcified sediments were then dried, weighed, and put into pre-cleaned tin cups for flash combustion. The stable isotope ratios are reported as δ values, in which δ=(R^sam^/R^std^–1)×1,000, with R being the isotope ratios (either ^13^C/^12^C or ^15^N/^14^N) in the sample and standard, respectively. The carbon isotope ratio of total organic carbon was referenced against Vienna Pee Dee Belemnite (VPDB). The nitrogen isotope ratios of total nitrogen were referenced against atmospheric N_2_. The analytical precision achieved through repeated analyses of in-house standards was typically better than 0.2‰ for both δ^13^C and δ^15^N.

### Cell counting

For determining total cell abundance in sediments collected during KR11-11, KR12-19 and KR14-01 cruises, microbial cells were fixed with sterile 0.2-µm-pre-filtered formaldehyde solution onboard and stored at −80°C until subsequent processes onshore as follows. The samples were centrifuged, washed with 4.5 mL PBS buffer and re-suspended in a PBS and 100% ethanol (1:1) mixture. After sonication using a Branson Sonifier 220 (Danbury, CT, USA), samples were diluted, filtered onto 0.2-μm polycarbonate filters, and stained using SYBR Green I. Excess stain was removed three times using 3 mL Milli-Q water, then filters were mounted onto microscope slides and cells were counted under blue light by epifluorescence microscopy Axioskop 2MOT (Carl Zeiss, Jena, Germany) at 1,000 × magnification. For sediments collected from the KR15-01 cruise, frozen samples were suspended in 5 mL PBS containing 4% formaldehyde and incubated at 4°C for 1–2 h. Samples were then centrifuged, repeatedly washed with PBS, re-suspended in 5 mL PBS and 100% ethanol (1:1) mixture, and stored at −20 °C. Fixed samples were diluted with PBS and sonicated for 20 s with an ultrasonic homogenizer UH-50 (SMT, Tokyo, Japan). Cells in the aliquant were stained directly using SYBR Green I and counted with an Olympus BX51 fluorescence microscope (Olympus, Tokyo, Japan) at 1,500 × magnification. For each filter, at least 400 cells were counted in at least 20 randomly chosen fields.

### DNA extraction, qPCR, and sequencing

Approximately 5 mL of frozen subsampled sediments from certain cores sections were used for DNA extraction. Environmental DNA was extracted using PowerSoil DNA Isolation Kit (Qiagen) with a minor modification to increase DNA yield: cells were shaken for 10 min after incubation at 65°C twice.

The abundance of prokaryotic and archaeal SSU rRNA genes was enumerated by qPCR analyses. Primer and probe sequences and PCR conditions are summarized in Table S2 [25, 26]. SSU rRNA gene copy numbers were quantified as averages of duplicates or triplicates for each sediment sample. For the qPCR analyses, 7500 Real-Time PCR System and StepOnePlus Real-Time PCR System (Applied Biosystems, Waltham, MA, USA) were used. For the preparation of qPCR mixtures, qPCR Quick GoldStar Mastermix Plus (Eurogentec, Seraing, Belgium) and Premix Ex Taq (Perfect Real Time) (Takara Bio, Shiga, Japan) were applied.

For SSU rRNA gene tag sequencing, the V4–V5 region of SSU rRNA gene was amplified from the environmental DNA assemblages using a primer set of U530F and U907R and LA Taq DNA polymerase with GC buffer (Takara Bio) [27, 28]. Primer sets and PCR conditions are summarized in Table S3. For amplification, Illumina adaptor sequence (ACACTCTTTCCCTACACGACGCTCTTCCGATCT) and Illumina Multiplexing PCR Primer 2.0 sequence (GTGACTGGAGTTCAGACGTGTGCTCTTCCGATCT) were added to the 5’ ends of the primers U530F and U907R, respectively. The amplicons were mixed with Illumina PhiX control libraries and sequenced by Illumina MiSeq platform (Illumina, San Diego, CA, USA) at JAMSTEC. The letter, serial number, ‘S’, and digits for each sample name represent the sampling site, experimental replication (if conducted), sample type (S for sediment), and upper sediment depth of subsampled layer.

### Bioinformatics

For raw sequence data, both ends of reads that contained low-quality bases (Phread quality score <20) and the adapter sequences were trimmed using TrimGalore (https://github.com/FelixKrueger/TrimGalore) with default settings. The remaining pair-end reads were merged with at least 10 bp overlap using FLASH [29] under default settings. Sequencing reads derived from PhiX genome were removed using Bowtie2 [30] using default settings. Chimeric sequences were filtered out using UCHIME [31] with default settings, and sequences with low complexity or shorter than 100 bp were discarded using PRINSEQ with *-lc_threshold 7* setting. All remaining high-quality sequences were clustered with a 97% identity threshold using VSEARCH [32]. After discarding singletons [33], each cluster was designated as an operational taxonomic unit (OTU). Rarefaction curves were calculated using RTK [34] with default settings. Non-metric multidimensional scaling (NMDS) analysis was conducted using Bray-Curtis dissimilarities, and multi-response permutation procedures (MRPP) test was conducted with 999 permutations and same dissimilarity indices. The taxonomic names of each OTU were systematically assigned using blastn search [35] against SILVA database release 132 [36] and retrieving the top hit sequence that showed e-values ≤1E-15.

A network structure of OTU co-occurrence was visualized using naive statistical metrics with strict cut-off values to consider a valid co-occurrence event. We used OTUs that represent >0.1% of the total sequencing pool, and pairwise correlations with >0.8 Spearman’s correlation coefficient (ρ) with Q-values <0.01 after Bonferroni correction; networks involved in more than two OTUs were analyzed.

SSU rRNA gene sequences of Thaumarchaeota were retrieved from the SILVA database [36] and systematically reduced in family level via similarity-based clustering with 90% identity using VSEARCH [32]. A maximum-likelihood (ML) tree was constructed using FastTree2 [37] with GTR+CAT option. Phylogenetic clades of Thaumarchaeota were affiliated following past studies [38, 39].

### Data deposition

Amplicon sequence data were deposited into the DDBJ Sequence Read Archive (SRA) under DRA008185 and DRA008316 (Table S1). All data were registered under BioProject ID PSUB010125.

## Results and discussions

### Organic geochemistry

Total organic carbon (TOC) and total nitrogen (TN) in the studied sediments ranged from 0.10 to 3.28 and 0.02 to 0.42 wt%, respectively (Fig. S1 and S2). Among the trench bottom sites, the highest TOC and TN values were detected in the Japan Trench sites (1.63–3.28 and 0.22–0.42 wt%, respectively) and the lowest were observed in the Mariana Trench sites (0.16–0.59 and 0.02–0.08 wt%, respectively), which differed by an order of magnitude. TOC and TN in the abyssal sediments slightly decreased with sediment depth, while those in the hadal sediments presented variable but decreasing trends. The C/N ratio was concordant with past observations that concentrations of protein, carbohydrate, and lipid were generally higher in the Izu-Ogasawara Trench region compared to the Mariana Trench (Manea et al. under review). The δ^15^N values of surface sediments may reflect nutrient availability at the ocean surface due to isotopic fractionation during nutrient consumption by phytoplankton [40]; nutrient-rich conditions can lead to lower values (5.1–5.4‰ and 2.4–8.2‰ at the Japan and Izu-Ogasawara Trenches, respectively), whereas nutrient-poor conditions cause higher values (8.9–11.7‰ at the Mariana Trench). Thus, the sedimental OM concentrations and traits likely reflect the different geographical settings and productivity of the investigated station. It is generally expected that the organic carbon deposition and subsequent diagenetic process reflects the surface productivity of the overlying ocean. The Japan Trench (station JC) is located under the relatively eutrophic north-western Pacific Ocean, where nutrient-rich Oyashio currents encounter warm Kuroshio currents; additionally, the close-distance to Honshu island, Japan, may contribute terrestrial OM to the seafloor [41, 42]. In contrast, the other stations are located under the oligotrophic Pacific Ocean, far from continents or large islands. In particular, the Mariana Trench region is located near a subtropical gyre known to have one of the lowest surface ocean productivities [43]. Even if definitive conclusions could not be made from the available data, the differences in C/N ratios of each site could reflect differences in OM sources as well as stable isotopic signatures.

### Pore-water chemistry

We measured concentrations of dissolved oxygen (O_2_) and pore water nutrients (NO_3_ ^−^, NO_2_ ^−^, NH_4_ ^+^, and PO_4_) in the obtained cores (Fig. 2 and S3) to study microbial decomposition of sedimentary OM using oxygen and/or nitrate as electron acceptors [44]. O_2_ concentrations decreased rapidly with increasing sediment depth in most sediment cores and depleted above 30 cm below seafloor (cmbsf), whilst those of abyssal cores collected from the Mariana Trench showed moderate decreases. In all hadal cores, NO_3_ ^−^ concentrations drastically decreased with sediment depth to less than 5 μM above 30 cmbsf with a concomitant increase in NH_4_ ^+^ concentrations, especially at the Japan and Izu-Ogasawara Trenches. In contrast, no apparent depletion of NO_3_ ^−^ (>27 μM) and lower NH_4_ ^+^ concentrations (<11 μM) were observed throughout the sediment depths in all abyssal stations from the Izu-Ogasawara and Mariana Trench regions, which also exhibited low TOC concentrations. Notably, NO_3_ ^-^ profiles showed large variations among hadal sediments compared to abyssal sediments. Among the NO_2_^−^ profiles, clear subsurface peaks up to 6.6 μM were found in only three sediment cores (3, 7, and 7 cmbsf of cores from JC, IO1-1 and IO1-2, respectively). PO_4_ concentrations generally increased with sediment depth, except for stations IO1-1 and IO1-2. These profiles of dissolved oxygen and nitrogen compounds are concordant with previous studies [4, 45]. The profiles suggested that microbial nitrate respiration was relatively modest in abyssal sediments down to 50 cmbsf, whereas the respiration was active in hadal sediments above 30 cmbsf, probably reflecting higher concentrations of fresh OM. Interestingly, the increase rate of NH_4_ ^+^ in hadal sediments along with sediment depth were different between northern and southern stations, suggestive of variance among stations in microbial populations and functions involved in nitrogen cycles.

**Fig. 2.**
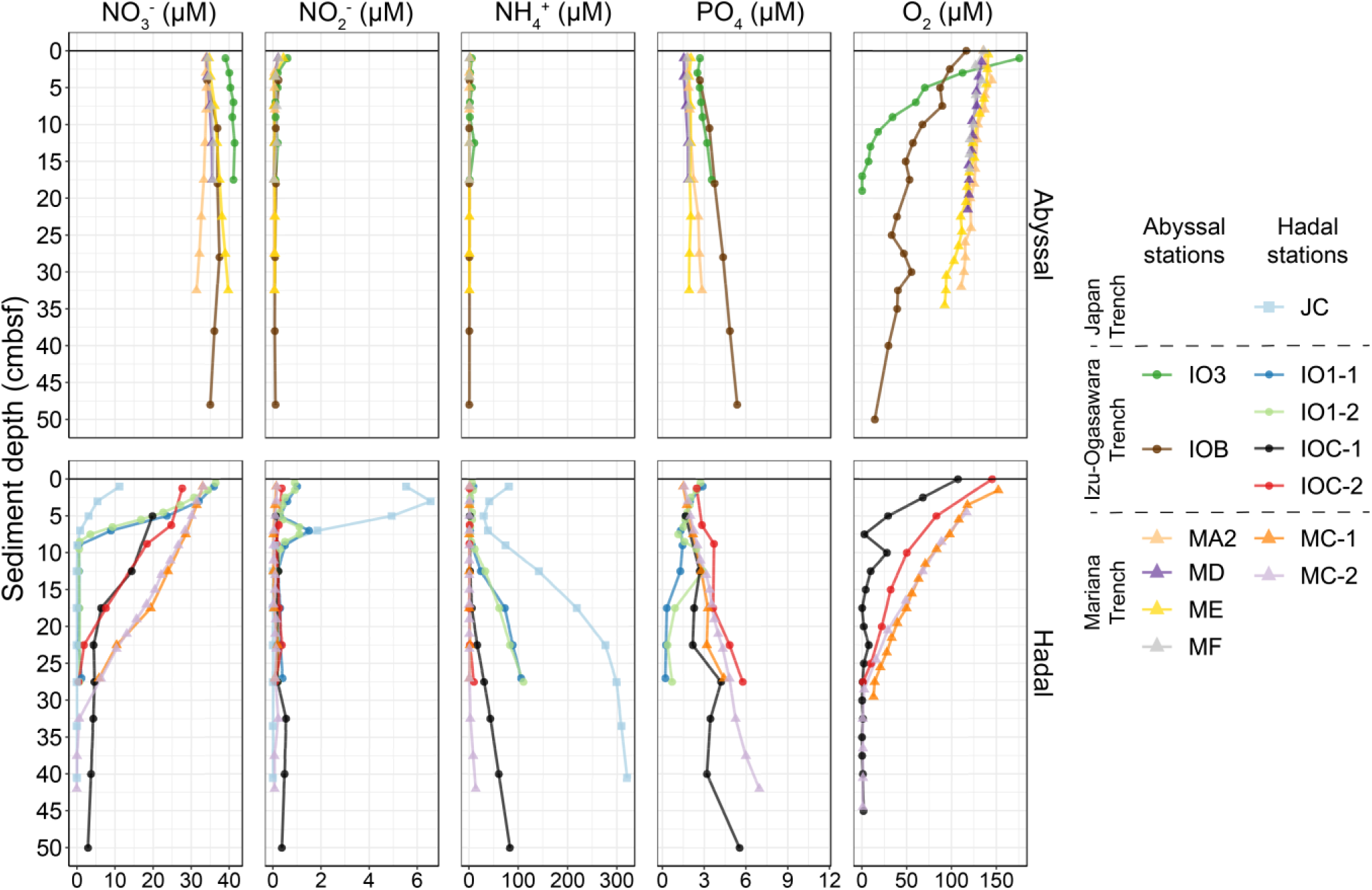
Porewater chemistry of the surface sediments in abyssal (upper panels) and hadal (lower panels) stations. Data down to 50 cmbsf are presented in this figure and entire sediment data are presented in Fig. S3.

### Microbial abundances

The direct cell counting and qPCR technique showed similar trends in microbial abundance (Fig. 3 and S4). Cell abundances by cell counting ranged from 4.2×10^5^ to 9.6×10^7^ cells/mL sediment, with a general decrease with sediment depth, while scattered profiles were found in the trench bottom sites of the Japan and Izu-Ogasawara Trenches (Fig. 3a and S4a). Cell abundances in the hadal stations were generally higher than those in the adjacent abyssal stations. When comparing maximum cell abundances between cores, the largest and smallest cell abundances in hadal sites were found in the Japan and Mariana Trenches, respectively; for abyssal sites, abundance at the Izu-Ogasawara Trench was larger than that of the Mariana Trench. For qPCR analysis, the maximum and minimum copy numbers of prokaryotic and archaeal SSU rRNA gene in each station ranged from 3.4×10^5^ to 3.0×10^9^ and 9.9×10^4^ to 5.7×10^8^ copies/mL sediment, respectively (Fig. 3b and S4b). Additionally, in the Mariana Trench region only, copy numbers from the shallow abyssal sediments were higher than those from the hadal sediments. The SSU rRNA gene copy numbers were 2–197 fold higher than the cell abundances by direct counting method, likely resulting from biases of primers, probes, extracellular DNA, and/or SSU rRNA gene copy numbers on prokaryotic genomes [46, 47]. However, cell abundance and SSU rRNA gene copy number was significantly correlated (Fig. S5). Interestingly, ratio of archaea rapidly depressed at approximately under 20 cmbsf of the hadal cores, while higher values were observed through the vertical profile in abyssal cores (Fig. 3c and S4c). Overall, these trends were consistent with previous studies [4, 48, 49], supporting more vigorous microbial activity in trench bottom hadal sediments, especially in the subsurface under 5 cmbsf.

**Fig. 3.**
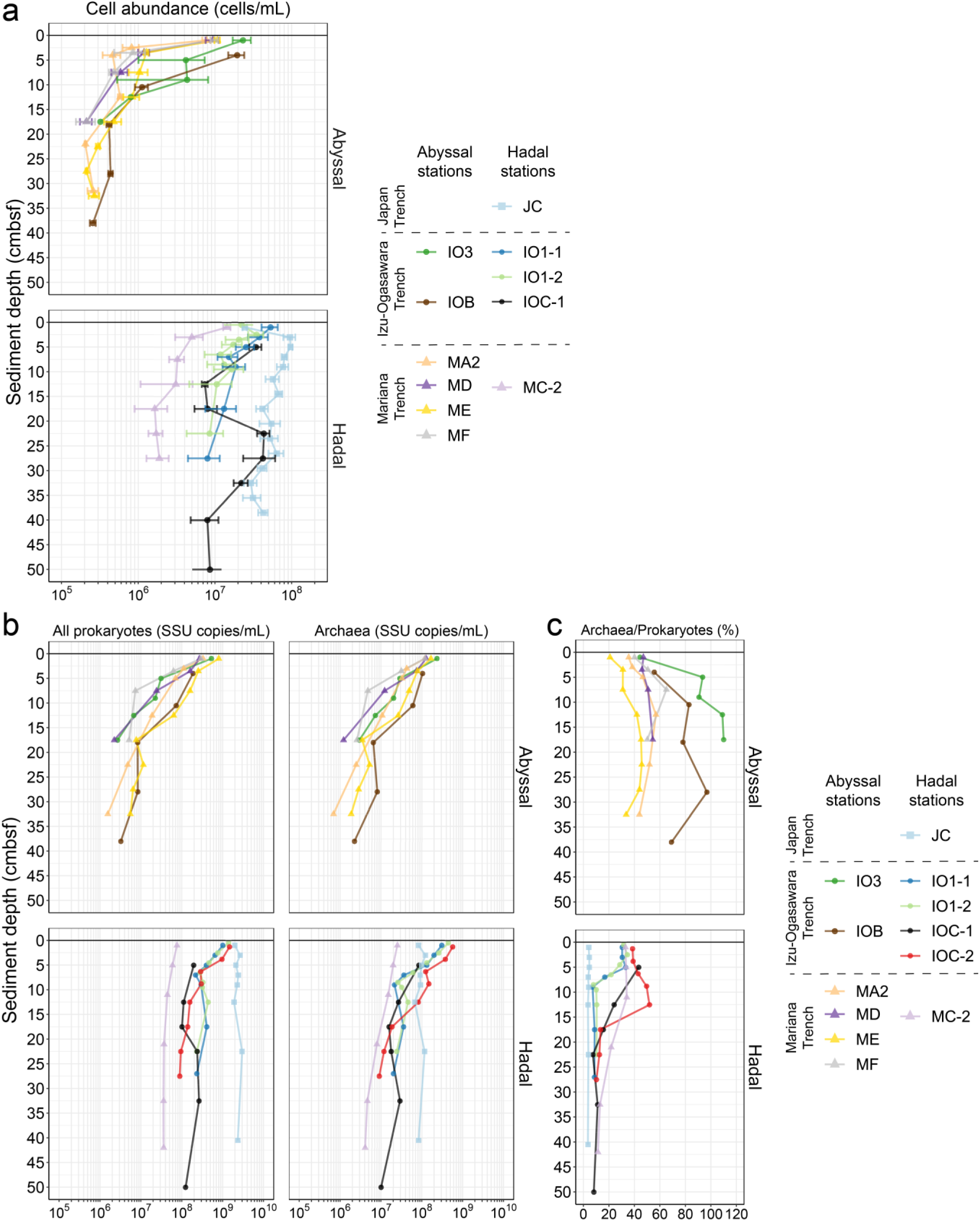
The abundance of microbes and ratios of archaea in each sediment core from abyssal (upper panels) and hadal (lower panels) stations. Abundances were measured using (**a**) cell counting and (**b**) qPCR techniques. The X-axis represents cell counts and SSU rRNA gene copies per milliliter of sediment, respectively. The error bars represent standard deviation. (**c**) Ratios of archaea/prokaryotes were calculated using qPCR data. Data from layers ranging between 0 to 50 cmbsf are presented in this figure, and full data are shown in Fig. S4.

### OTU-level compositions of microbial communities in sediment samples

Based on the geochemical profiles in sediments, especially dissolved oxygen and nitrate concentrations, we selected four to ten layers from each sediment core for SSU rRNA gene tag sequencing. A total of 8,286,508 high-quality SSU rRNA gene sequences with 414 bp average length were obtained from the 92 sediment subsamples. The sequences comprised of 80,478 OTUs with 1,587–13,181 (5,478 average) OTUs per sample (Table S1). Based on rarefaction curves, the obtained OTUs in each sediment sample well-represented their microbial communities (Fig. S6).

To investigate compositional similarity between samples, we performed OTU-based NMDS analysis. The OTU compositions were significantly structured along the stations (A = 0.31, p<0.001, MRPP) (Fig. 4a) and differed between the abyssal and hadal sediments (Fig. 4b). Significant associations were also observed with each of the three trenches, especially among the hadal cores (A = 0.21, p< 0.001, MRPP). Unexpectedly, the compositions in abyssal sediment from the Izu-Ogasawara and Mariana Trench regions overlapped. The separation among the abyssal sediments from the Izu-Ogasawara and Mariana Trench regions correlated with the different depression trends in porewater nitrate, TOC, and TN concentrations, but not with porewater oxygen concentration and cell abundance (Fig. 2, 3, and 4c). OTUs were generally shared along with sediment cores at each station (Fig. 4d).

**Fig. 4.**
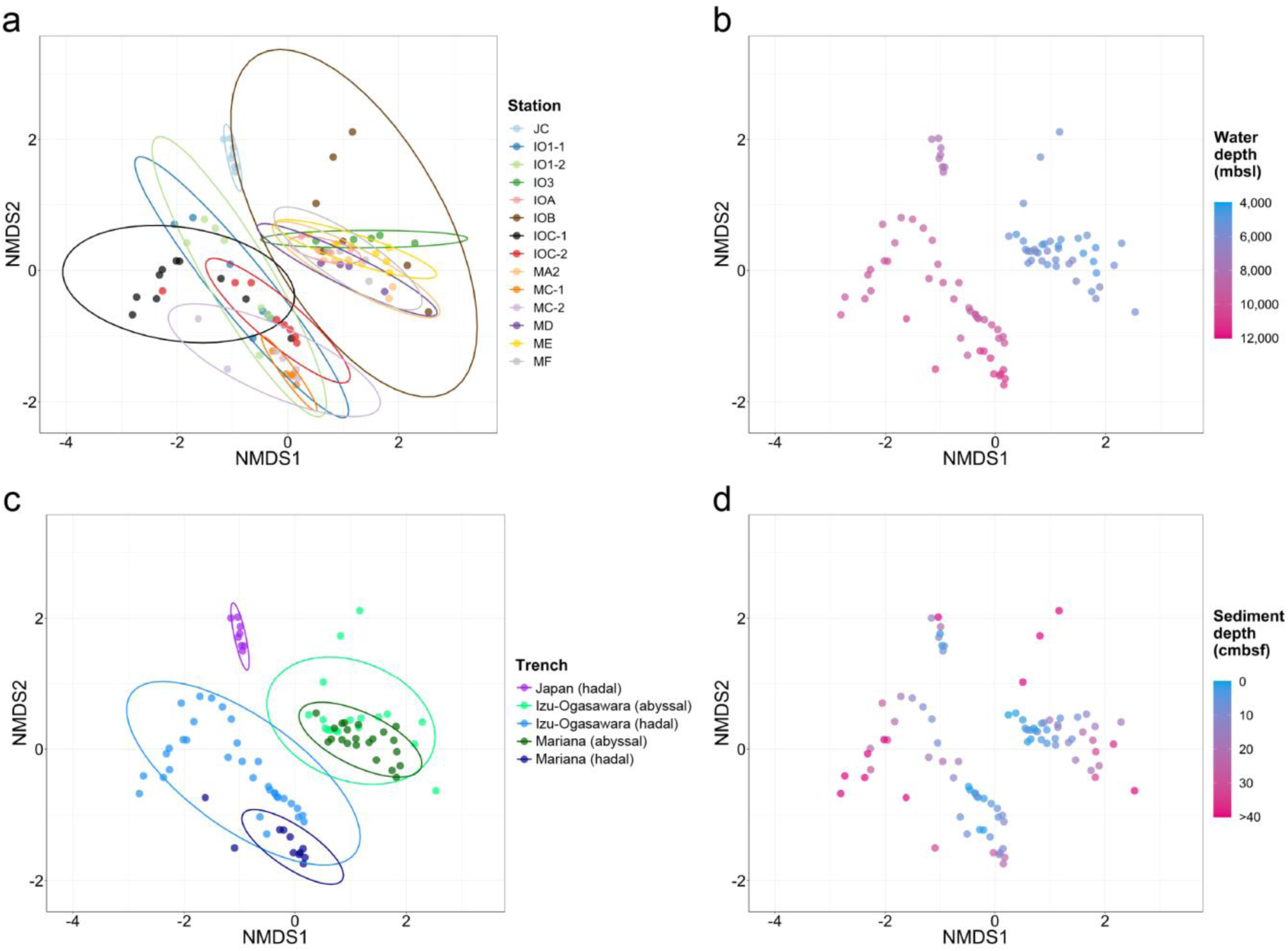
Nonmetric multidimensional scaling (NMDS) plots for OTU compositions. The distance matrix was calculated based on the Bray-Curtis dissimilarity. The stress value of the final configuration was 18.2%. The sediment samples were colored depending on (**a**) sampling station, (**b**) water depth, (**c**) trench and geomorphology, and (**d**) sediment depth. (**a**,**c**) Ellipsoids represent a 95% confidence interval surrounding each group, and each sediment core and geography are coded by color.

Vertical profiles of community diversity in the hadal sediment cores were distinct from those of the abyssal sediment cores (Fig. S7). The Shannon index values of all abyssal sediment cores gradually decreased with sediment depth, while those in hadal samples fluctuated. Overall, the values in the upper most layers at the abyssal stations were higher than those at the hadal stations. In hadal sediment cores, index values were depressed at approximately 5–20 cmbsf and retained or increased below these layers. In all hadal cores from the Izu-Ogasawara Trench, clear peaks were found at 8–25 cmbsf and the values were higher than those of the most surface layers. Conversely, in cores from the Mariana Trench, we observed slightly increasing tendencies in deep sediment sections, indicating that peaks were possibly located in layers deeper than 50 cmbsf. The diversity-depression was observed close to layers where depletion of oxygen and nitrate occurred (Fig. 2). These trends were concordant with the scattered microbial cell abundances observed in most of the hadal sediments in contrast to the abyssal sediments (Fig. 3), but opposed to the general trends that prokaryotic growth and bioactivity are restricted according to sediment depth in energy-limited subseafloor sediments [42, 50, 51], which should be an unique feature of hadal subsurface biosphere. In the cases of gut and freshwater environments, correlation of the supply of fresh nutrient resources and microbial biomass and diversity has been discussed [52, 53]; thus, the feature of the hadal subsurface biosphere could be explained by the deposition of relatively fresh organic compounds with high sedimentation rate in the hadal seafloor.

### Taxonomic composition of microbial communities in sediment samples

Among the retrieved OTUs, 76,881 (99.7%) were taxonomically assigned to eleven and two phyla in Bacteria and Archaea, respectively. Only 153 OTUs were assigned to Eukarya and the remaining 243 OTUs were taxonomically unassigned. The top three and ten most abundant phyla accounted for >58% and >88%, respectively, of the sequence pool of all sediment samples.

Proteobacteria (average 23.8%) was the most abundant phylum, followed by Thaumarchaeota (23.7%), Planctomycetes (10.6%), and Chloroflexi (9.6%) (Fig. 5). Within sequencing reads assigned to Nanoarchaeaeota, 99.6% were assigned to class Woesearchaeia, currently proposed as novel phylum *Candidatus* (*Ca.*) Woesearchaeota. Thus, we assigned Nanoarchaeaeota as Woesearchaeota in this study. Within Proteobacteria, Gammaproteobacteria (10.3%) is the most abundant class, followed by Alphaproteobacteria (8.4%) and Deltaproteobacteria (5.1%) (Fig. S8). In general, the dominant phyla were similar to those of previous studies of abyssal and hadal sediments [20, 21, 48, 54–56].

**Fig. 5.**
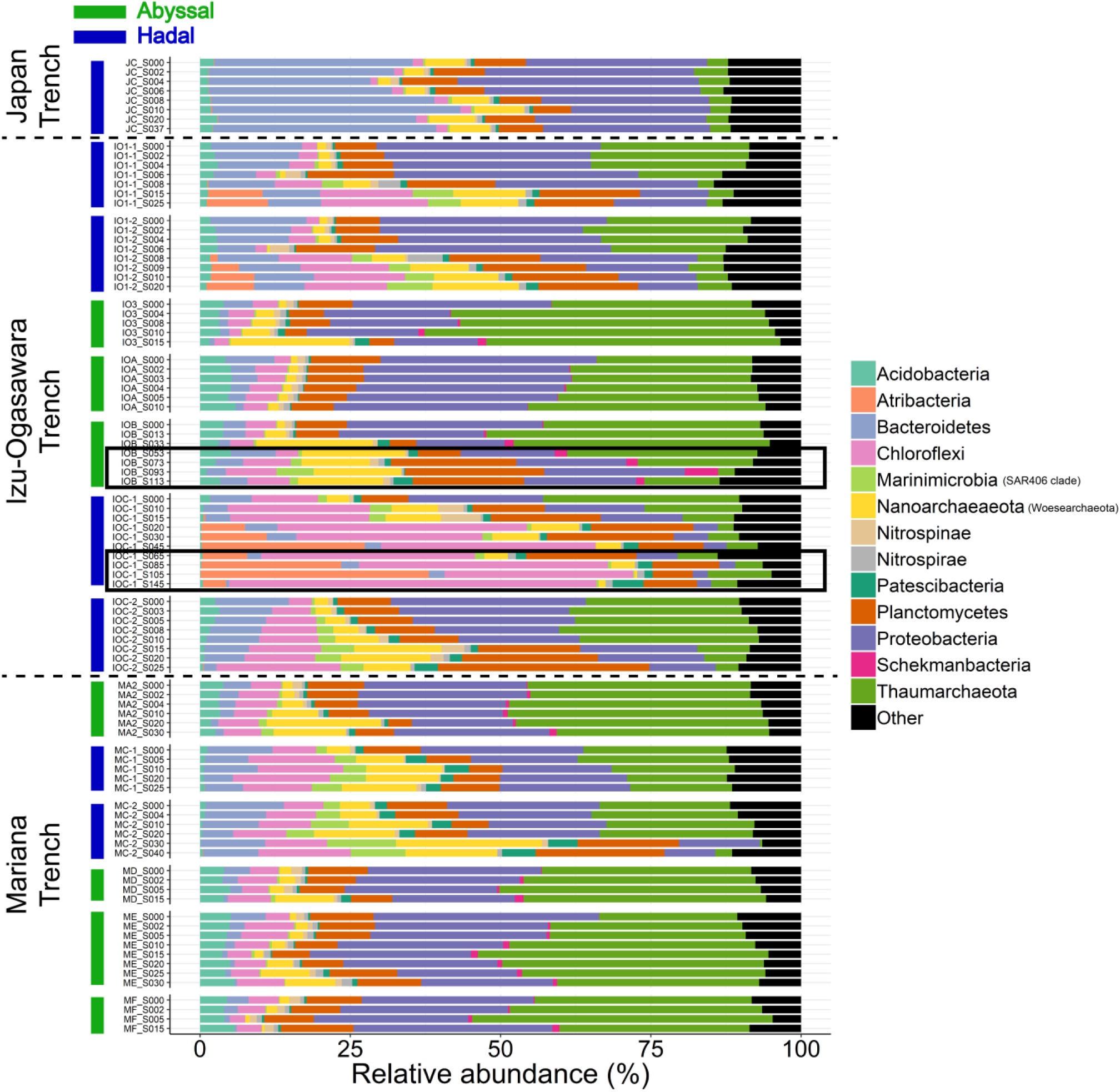
Relative abundances of sequences at the phylum level. Groups with <5% abundance were summarized as ‘Other’. Sediment samples retrieved from deep sediment layers (>50 cmbsf) are indicated by surrounding black rectangles.

Although dominant phyla were shared among subsamples within each of the sediment cores, their relative abundance gradually changed with sediment depth (Fig. 5). The relative abundances of Chloroflexi, Woesearchaeota, and *Ca.* Marinimicrobia generally increased in deeper sections along with the gradual decreases in oxygen and nitrate concentrations (Fig. 2). Conversely, Proteobacteria and Thaumarchaeota dominated in shallower sections. These trends are similar to previous studies [20, 21, 48, 54–56].and likely depend on the concentrations of dissolved oxygen and nitrate in sediments. Woesearchaeota has been detected from diverse benthic and anaerobic environments [57–61] and harbors genomic capability for fermentation-based metabolism [62]; hence, they may contribute to anaerobic carbon and hydrogen cycles in the deep seafloor sediments.

We also observed some notable differences in prokaryotic communities likely associated with geographical location. Distinct community compositions were observed in station JC; Bacteroidetes predominated the communities (average 33.7%), while Thaumarchaeota was relatively scarce (4.1%). Within the Bacteroidetes, Flavobacteriaceae (belonging to class Flavobacteriia) is the most abundant family. Because members of Flavobacteriia were reported to be abundant in eutrophic oceans [63], our results likely reflect eutrophic productivity in the Japan Trench. The predominance of phyla *Ca.* Atribacteria was found only in deeper sections of IO1-1, IO1-2, and IOC-1 hadal stations at the Izu-Ogasawara Trench. Atribacteria is a common lineage in organic rich anaerobic sediments and probably grow with fermentation [64–66]. The higher abundances of Flavobacteriia and Atribacteria may represent a substantial deposition of degradable organic compounds into hadal sediments.

The substantial differences revealed in the taxonomic analysis may also be explained by the geomorphological variations among sampling stations. Marinimicrobia showed significant unevenness between cores, with higher abundances observed in hadal (2.6%) verses abyssal sediments (0.73%) (p<0.05, U-test, Bonferroni correction). Marinimicrobia is known to be a dominant population in deep sea sediments and seawater, especially in oxygen-minimum zones, and harbors genetic potential of diverse energy metabolic processes using sulfur and nitrogenous compounds as electron donor and acceptor [67–71]. In contrast, *Ca.* Schekmanbacteria was detected in all abyssal samples (average 0.82%), while there was significantly lower abundance (0.007%) in hadal samples (p<0.05, U-test, Bonferroni correction). Since these phyla were recently proposed [72], their biological and geochemical importance remains unclear.

The relative abundance of Thaumarchaeota showed drastic decrease in hadal stations above 6-15 and 20–30 cmbsf in core(s) from the Izu-Ogasawara (IO1-1, IO1-2, IOC-1, and IOC-2) and Mariana (MC-2) Trenches, respectively, where nitrate was consumed and ammonium emerged (Fig. 2). In contrast, low abundance of Thaumarchaeota was continuously observed in sediments from the Japan Trench (JC), where nitrate concentration was depleted through the sediment core (Fig. 2). This was concordant with qPCR analyses (Fig. 3c), as well as previous observations that Thaumarchaeota frequently dominated in aerobic sediment columns and radially decreased across the oxic-anoxic transition layer [55, 73]. The most predominant family of Thaumarchaeota in the sediments was Nitrosopumilaceae (92%), which are known to be ammonia oxidizing archaea (AOA) [74–76], and thus may contribute markedly to nitrification processes in trench surface sediments. The co-existence of Thaumarchaeota and Marinimicrobia at relatively high abundance suggests the co-existence of nitrification and denitrification processes, respectively, as described previously [48, 77]. Although abundance of functional genes related with nitrification (*e.g., amoA*) was not analyzed in this study, we should note that abundance of the *amoA* gene decreased with sediment depth in trench bottom sediments from the Mariana and Izu-Ogasawara Trenches [48, 77].

### Co-occurrence network structure of OTUs

Many prokaryotic lineages are known to establish consortiums with specific prokaryotic members, sharing similar ecological niches and biological interactions (e.g., sharing metabolic compounds via fixation and translocating process) [78]. Since co-occurrence patterns can be useful for revealing such concrete but mostly hidden relationships from complex community datasets, co-occurrence network analyses have been widely applied to various SSU rRNA tag sequencing datasets of marine and other environments [79–82]. Here, we conducted co-occurrence analysis to understand core metabolic interactions among microbes in trench subseafloor sediments.

The co-occurrence network showed six clusters composed of 3–36 OTUs and 2–247 edges, and a total of 66 OTUs represented average 23.6% in each sample (Fig. 6). Interestingly, most prokaryotic alliances were likely related to oceanographic zonation (i.e., abyssal and hadal) and sediment depth (Fig. 7). OTUs belonging to the largest group A (composed of two subgroups A-1 and A-2) were abundantly detected in the abyssal sediments through the Izu-Ogasawara and Mariana trenches, whilst abundance OTUs in other groups were higher in hadal sediments: under 10 cmbsf for group B, in shallow sections (0–30 cmbsf) for group C, and above 6 cmbsf for group D. The members of groups E and F were detected from both abyssal and hadal sediments cores. Although OTUs of group C and F were spread among the three trenches, those of group B, D, and E were rare in cores from the Japan Trench.

**Fig. 6.**
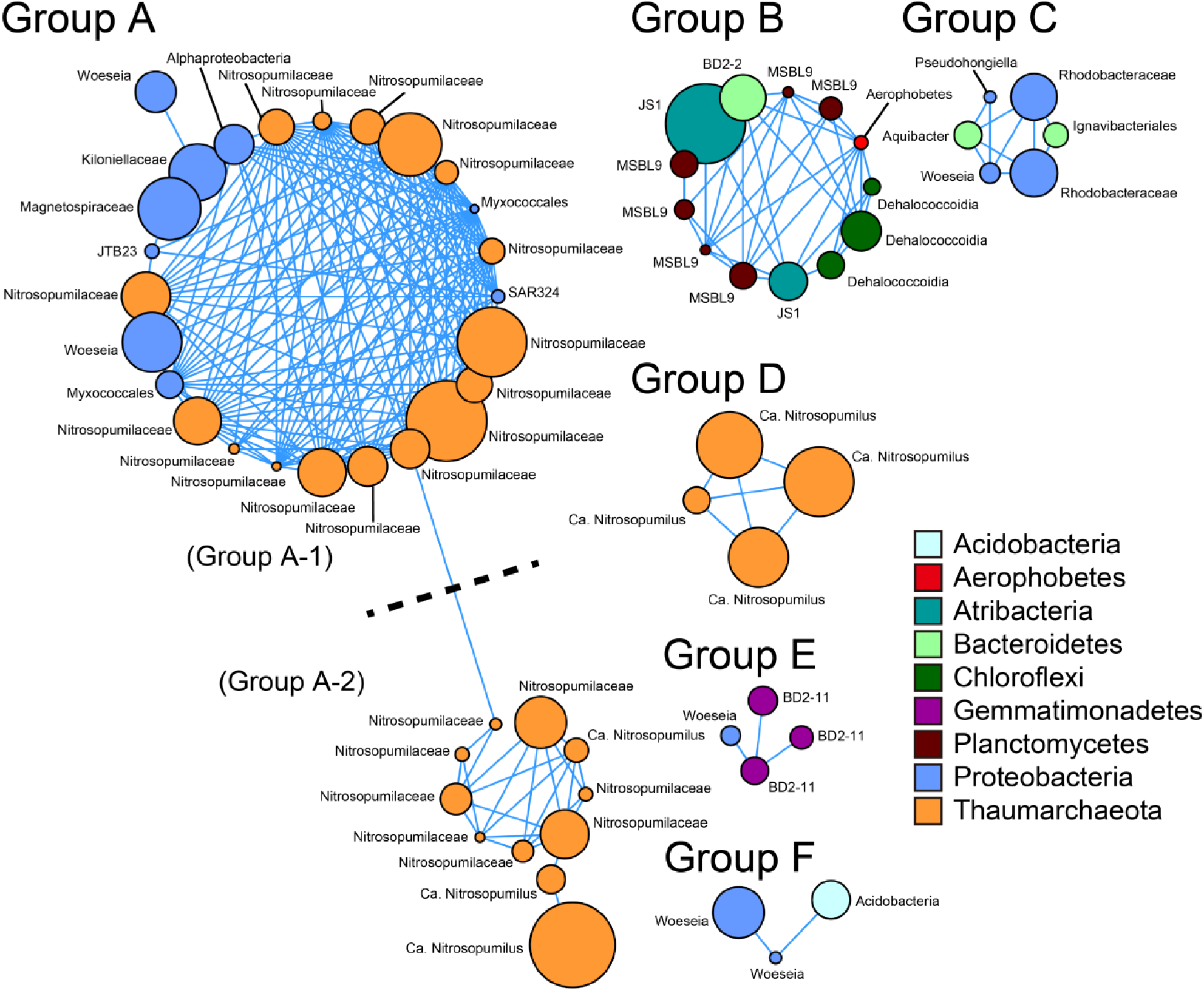
Architecture of co-occurrence OTU networks. Nodes represent OTUs and edges represent statistically significant positive correlations of each OTU pair. The size of nodes represents relative abundance of OTUs in the data set. Nodes are colored by taxonomy at the phylum level.

**Fig. 7.**
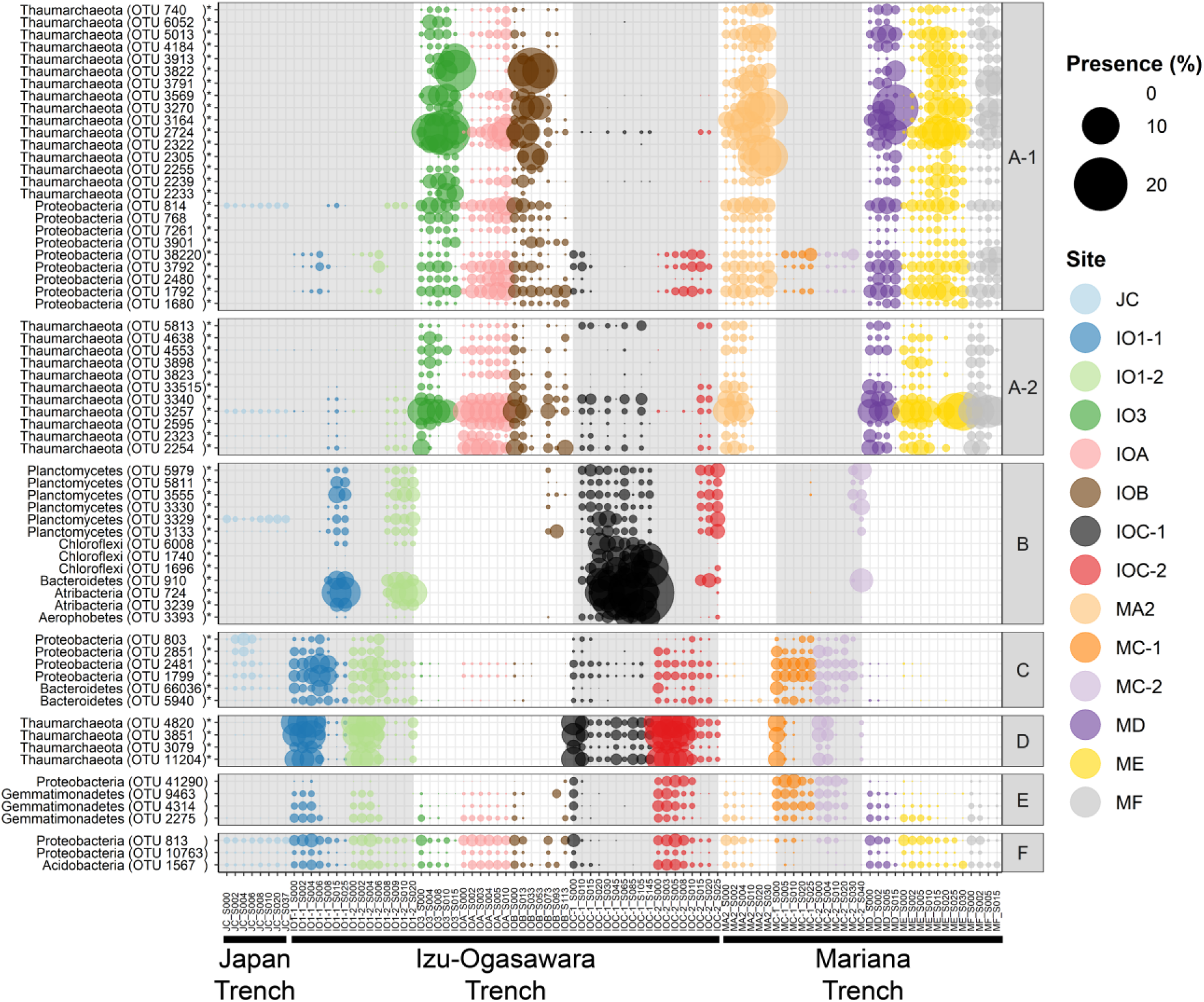
Bubble plot showing comparative OTU profiles belonging to each co-occurrence network. Bubbles are colored by sampling station and bubble sizes correspond to relative abundances. The white and gray backgrounds represent abyssal and hadal sediment samples, respectively. A to F written in the right side represent co-occurrence network groups summarized in Fig. 6. OTUs with asterisks indicate statistical significance of localization in either the abyssal or the hadal samples (p<0.05, U-test, Bonferroni correction).

Groups A and D were dominated by OTUs assigned to order Nitrosopumilales (Fig. S9). Nitrosopumilales are considered as aerobic nitrifying [74–76], and abundance of these OTUs were concordant with the high oxygen and nitrate concentrations in the sediments (Fig. 2). OTUs of group D belonged to one small clade (showing 95.6–96.1% identity with 16S rRNA gene of *Nitrosopumilus maritimus* SCM1 [DQ085097]) while those of group A were spread throughout the order. The niche separation of AOA subgroups is regulated by the availability of electron donors like ammonia [19, 83, 84]. Supporting this, the dominance of the group D clade (up to 78.1% in Thaumarchaeota in the hadal samples) is consistent with the enrichment of labile OM in the hadal sediment habitats. Unfortunately, further subgroup assignment of Thaumarchaeota OTUs using short 16S rRNA gene tag sequences were not technically feasible in contrast to the previously described *amoA* gene [38]. Although most proteobacterial OTUs in group A were assigned to lineages that currently less well-understood, two OTUs were assigned to genus *Woeseia* in Chromatiales; this genus likely possesses denitrification pathway-related genes [85], and may consume nitrates that provided by nitrifiers including Thaumarchaeota.

Intriguingly, group B consisted of 13 OTUs belonging to five phyla (Planctomycetes, Atribacteria, Chloroflexi, Bacteroidetes, and *Ca.* Aerophobetes), and OTUs were abundant in the deeper sections where oxygen and nitrate were depleted (Fig. 2). All 13 OTUs showed highest sequence similarities with uncultured lineages, most of which were found in anaerobic environments with low oxygen concentrations. Atribacteria partake in fermentation metabolisms that produce acetate, hydrogen and carbon dioxide [64–66]. Similar to Atribacteria, recently defined bacterial phylum Aerophobetes were reported to harbor saccharolytic and fermentative metabolism capabilities [86]. All Planctomycetes OTUs were assigned to order MSBL9. Metagenome sequencing analyses identified genes encoding pyruvate formate-lyase from a member of MSBL9 [87], indicative of fermentation capabilities. All three Chloroflexi OTUs were assigned to class Dehalococcoidia, which is frequently detected from seafloor sediments and possesses genes related to hydrogen and sulfur compound oxidation with reductive dehalogenation of halogenated organic compounds [88–90]. Notably, MSBL9 and Dehalococcoidia possess potential of flavin secretion in marine sediments [91], implying that these lineages also contribute to maintaining extracellular metabolite pools in hadal sediments. The single Bacteroidetes OTU was affiliated with class BD2-2, which may interact with methanotrophic archaea and sulfate-reducing bacteria in methane seep sediments [92]. While knowledge of group B OTUs is still limited, they may cooperatively establish anaerobic metabolic networks; e.g. products of fermentation by Atribacteria, Aerophobetes, and MSBL9 are used as electron donors by BD2-2 and Dehalococcoidia OTUs, and then the fresh labile OM will be used as energy resources again in hadal sediments following necromass turnover recycling, as discussed previously [93].

Overall, the network structure analyses highlighted associations between prokaryotic consortiums and geochemical conditions (geomorphological zonation and sediment depth), with these consortiums likely representing potential metabolic interactions with habitat transition. In addition, the consortium structures were widespread among the trenches in the northwest Pacific Ocean. While three groups (B, C, and D) were selectively abundant in the hadal sediments, only one group (A) showed high preference in the abyssal sediments (Fig. 7).

### Factors impacting hadal subseafloor ecosystem

Here, we conducted culture-independent molecular analyses of trans-trench prokaryotic communities in the abyssal and hadal sediments collected from three different trench systems under different oceanographical settings to understand the general role of hadal environments on subsurface geochemical cycles and microbial ecosystems. Microbial cell abundance showed greater biomass in the hadal sediments verses abyssal sediments especially in deeper layers, which is consistent with previous studies [4, 48, 49]. Overall, the microbial composition suggested that development of prokaryotic communities depends on ocean geomorphological zonation (i.e., abyssal vs. hadal), geographic regionality (i.e., productivity of overlying ocean-surface), and factors associated with sediment depth. We also observed higher diversity of microbial communities among hadal trench sediments than abyssal plains and identified potential prokaryotic consortiums that spread their inter–trenches habitats and likely shared energy-retrieving metabolic processes in both abyssal and hadal sediments.

In general, deposition of OM and subsurface microbial cell abundance is related to productivity of the surface ocean and distance from continents or islands [42]. Indeed, in our abyssal sites, greater microbial cell abundance and oxygen consumption were observed in the Izu-Ogasawara compared to Mariana trench regions (Fig. 2 and 3). However, microbial compositions and some geochemical parameters (nitrate, TOC, and TN) were unexpectedly similar between these two regions, indicating that the impact of surface productivity on subsurface microbial community is much smaller than expected in the abyssal plains under oligotrophic to ultra-oligotrophic oceans. In the hadal sites, we found variations in geochemical parameters, cell abundances, and microbial compositions along with surface productivities. The differences between the abyssal and hadal sediments cannot only be explained by the vertical flux of sinking organic particles influenced by the ocean surface productivity. To explain the variations, we have two hypotheses: presence of intrinsic OM supplies on hadal sites apart from those sinking from the ocean surface, and impact of surface productivity on subsurface microbial community appears profoundly in hadal and modestly in abyssal sediments because of very high and low sedimentation rates, respectively.

Lateral transport along the trenches is one of the possible sources of OM in trench bottom sediments apart from sinking OM. There are north- and south-ward currents along the trench axis at the Izu-Ogasawara and Japan Trenches, respectively [94, 95]. These currents may contribute to transportation of suspended particles with relatively high OM contents from the north to south. A part of the latitudinal gradients in TOC values among hadal sites (Fig. S1) could be explained either by this lateral OM supply, or benthic microbial populations that prefer subsurface ecosystems under eutrophic oceans represented by Atribacteria (Fig. 5). However, we could not find clear geochemical signatures supporting OM delivery along the trenches.

OM degradation process at the surface sediments may differ between hadal and abyssal sites due to differences in sedimentation rates, which are very high at hadal while low at abyssal sites. Extremely high sedimentation rates at trench bottoms cause the rapid deposition of OM into the subsurface [9, 49]. This burial prevents oxidative degradation of OM at surface sediments, allowing semi-labile OM to be available to subsurface microbes. In contrast, abyssal plains generally have slower sedimentation rates. In our studied Izu-Ogasawara Trench sites, estimated sedimentation rates were 25 and 2.9 cm per 1,000 years at the hadal (IOC-2) and abyssal (IOA) stations, respectively, based on bulk ^14^C-age analysis (Nomaki et al., unpublished data). The slower sedimentation rates at abyssal sites have allowed continuous oxic degradation of OM at surface sediments for over 1,000s years, and labile OM are likely more diminished than those at hadal sites. Consequently, the differences in TOC concentrations (0.11–0.55%) were small across the abyssal plains in our sites, while those at the hadal trenches varied substantially (0.16–3.28%) (Fig. S1). The variations in TOC concentration among the hadal trench sediments subsequently influenced the dissolved oxygen and nitrate profiles through sediment depths (Fig. 2). Higher diversity of microbial communities among hadal trench sediments than abyssal plains (Fig. 4c) may reflect variations in TOC and porewater chemistries. Moreover, the differences in labile OM deposition may be reflected in the microbial cell abundance in the abyssal stations (Fig. 3) instead of microbial community structures.

Our findings provide new perspectives into hadal biospheres under different oceanographic regions, displaying contrasting properties to abyssal biospheres. We also identified novel insights into abyssal geochemistry and microbial communities whereby variations in surface productivity at abyssal sites are not profound in oligotrophic to ultraoligotrophic areas because the OM buried into subsurface sediments are extensively degraded before burial. However, we could not specify the factors controlling microbial ecosystems and biogeochemical cycles in this study. To further understand such controlling factors, investigation of microbiological processes in intra- and inter-cell scales and elucidation of biological mechanisms of trench systems that impact hadal biospheres are necessary. Ecological functions and phylogenetic classifications of most predominant lineages in deep seafloor sediments remain largely unknown. Moreover, although bacteria and archaea may account for a major part of ecosystems, viruses could facilitate biogeochemical cycles though biological interactions with prokaryotes in oligotrophic deep sea environment [93, 96–98]. Therefore, further taxonomic composition analyses, gene- and genome-centric approaches (e.g., metagenomics, metatranscriptomics, and metaepigenomics [99]) and integrative analyses with viruses (e.g., viromics) will provide further insights into microbial ecology and associated biogeochemical cycles.

## Contributions

SH conceived of and designed the study, performed the bioinformatics analyses, and wrote the manuscript. MH designed the experiments, sample collection, and performed the DNA extraction, qPCR analyses, and DNA sequencing. YM, AM, and MN designed and performed the geochemical experiments and analysis. MT performed the DNA extraction and qPCR analyses. J performed the DNA extraction, qPCR analyses, and cell counting. ER, RD, and CC performed the sample collection and cell counting. HM, TK, and ET contributed for the sediment sample collection. KT supervised the project. HN designed the geochemical experiments and analysis and wrote the manuscript. TN conceived of and designed the study, wrote the manuscript, and supervised the project. All authors read and approved the final manuscript.

## Acknowledgments

We would like to thank the captain, crew, and onboard scientists and technicians of the R/V *Kairei* (JAMSTEC) during KR11-11, KR12-19, KR14-01, and KR15-01 cruise, and the R/V *Yokosuka* (JAMSTEC) during YK11-06 cruise. We are very grateful to HOV *Shinkai 6500* and ROV *ABISMO* development and operation team. We thank Osamu Koide, Takaaki Kubota, Tomoko Makita, Masashi Tsuchiya, Kentaro Inoue, Mitsuhiro Yoshida, and Katsunori Yanagawa for sediment sampling, and Megumi Kuroiwa, Tomoko Makita, Shota Takino, Yasuhito Ito, Kazuki Shinoda, and Keisuke Koba for geochemical analyses. We also thank Shinsuke Kawagucci and Taichi Yokokawa for their helpful suggestions.

