## Supplemental Information for "Microbial community and geochemical analyses of trans-trench sediments for understanding the roles of hadal environments"

Figure S1 to S9.

Table S1 to S3.

#### Supplementary Figures

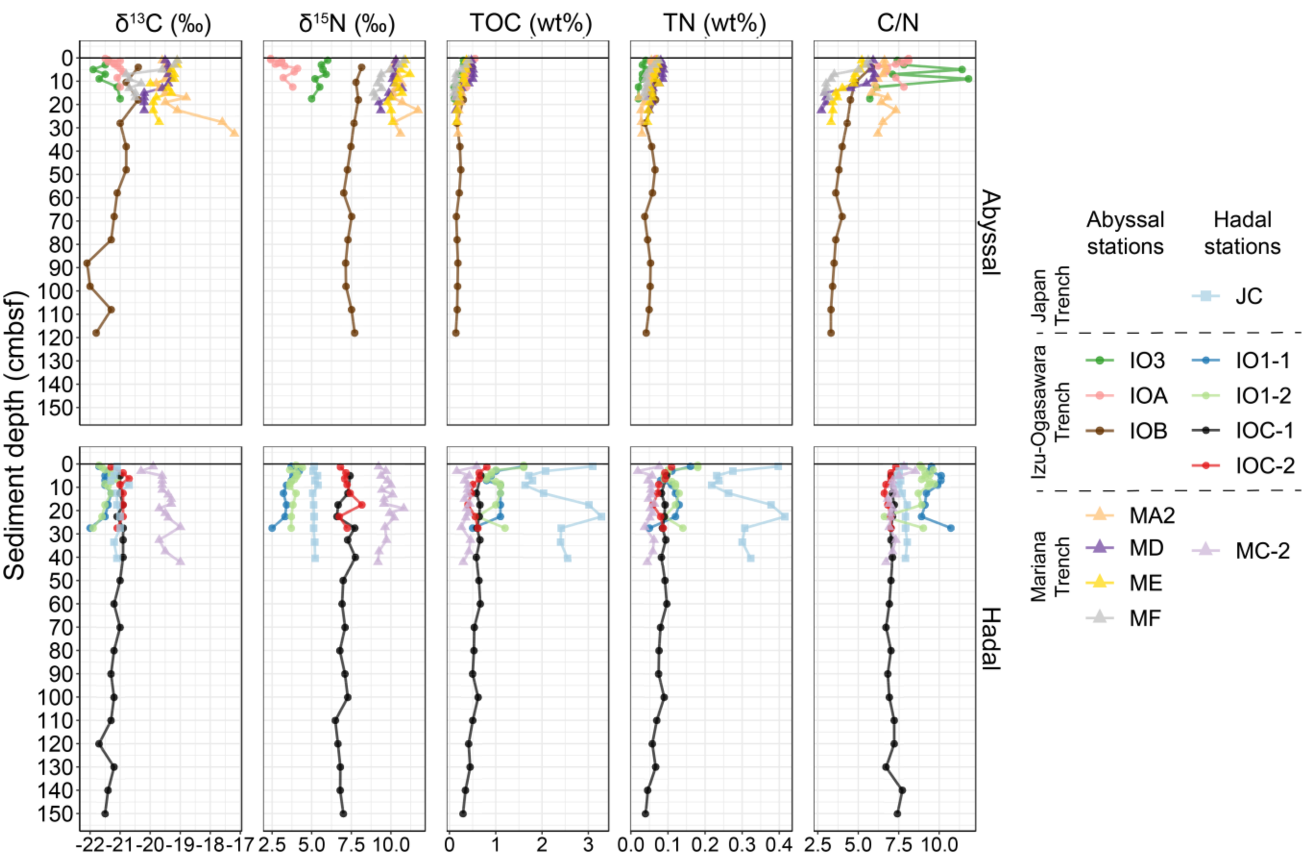


**Figure S1.** Concentrations of C and N isotopes ($\delta^{13}C$ and $\delta^{15}N$, respectively), total organic carbon (TOC), total nitrogen (TN), and C/N ratio of the surface sediments in abyssal (upper panels) and hadal (lower panels) stations.

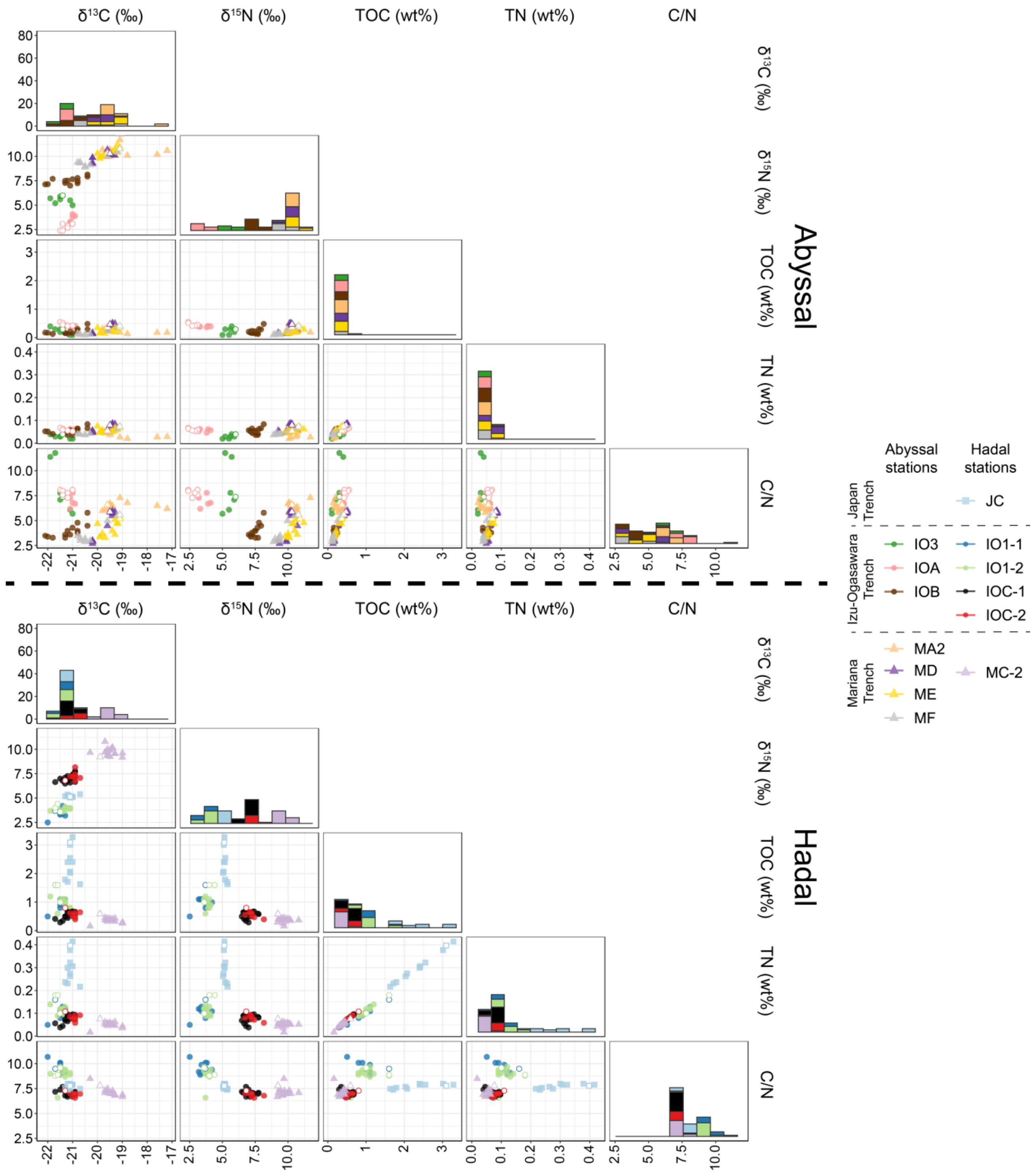


**Figure S2.** Pairwise relations of total organic carbon (TOC), total nitrogen (TN), C and N isotopes ($\delta^{13}$C and $\delta^{15}$N, respectively), and C/N ratio of the sediments in abyssal (upper panels) and hadal (lower panels) stations. The lower triangle panels indicate scatter plots of each pair of the geochemical measurements. Each sediment core is coded by color. Blank nodes represent measurements from surface layer (<3 cmbsf). The diagonal plots indicate cumulative histograms of measurements.

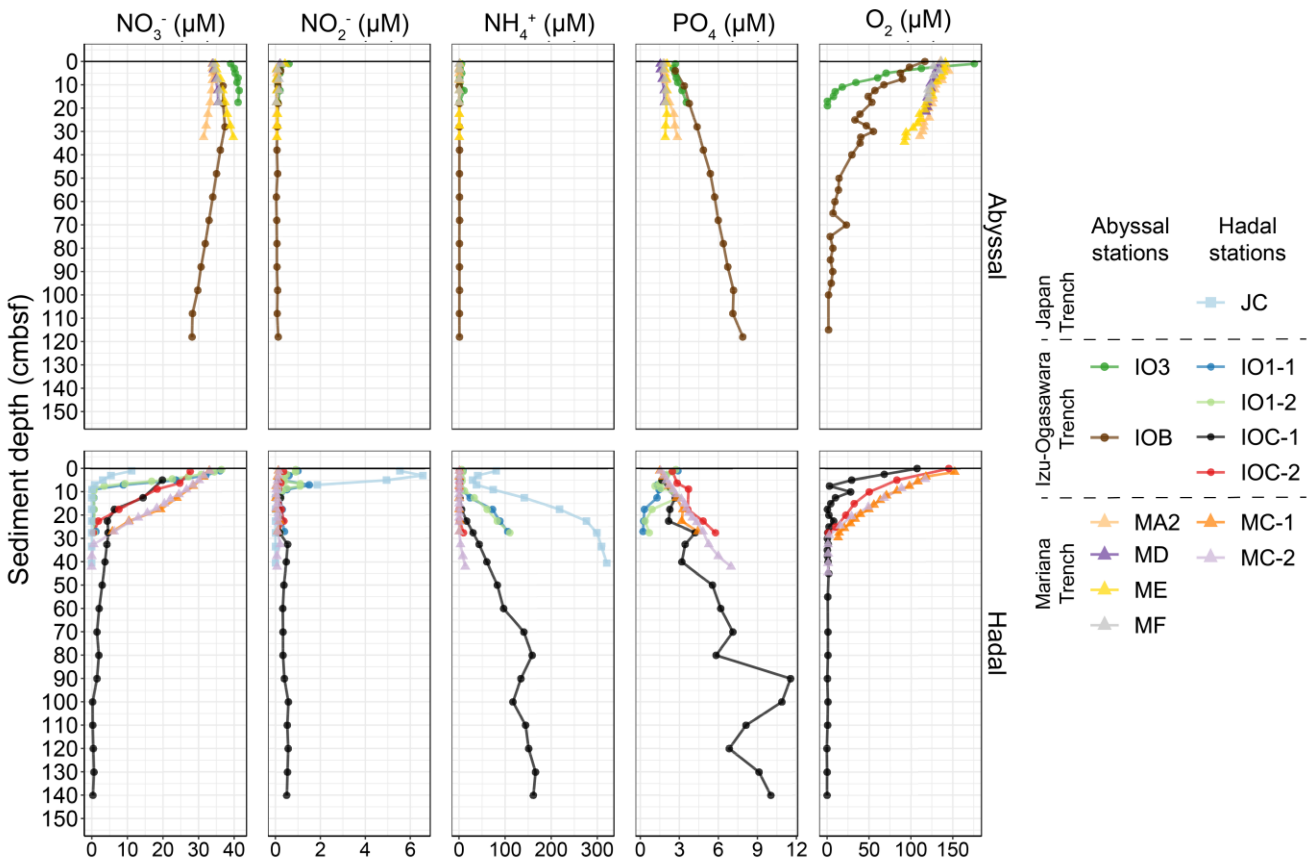


**Figure S3.** Porewater chemistry of the surface sediments in abyssal (upper panels) and hadal (lower panels) stations.

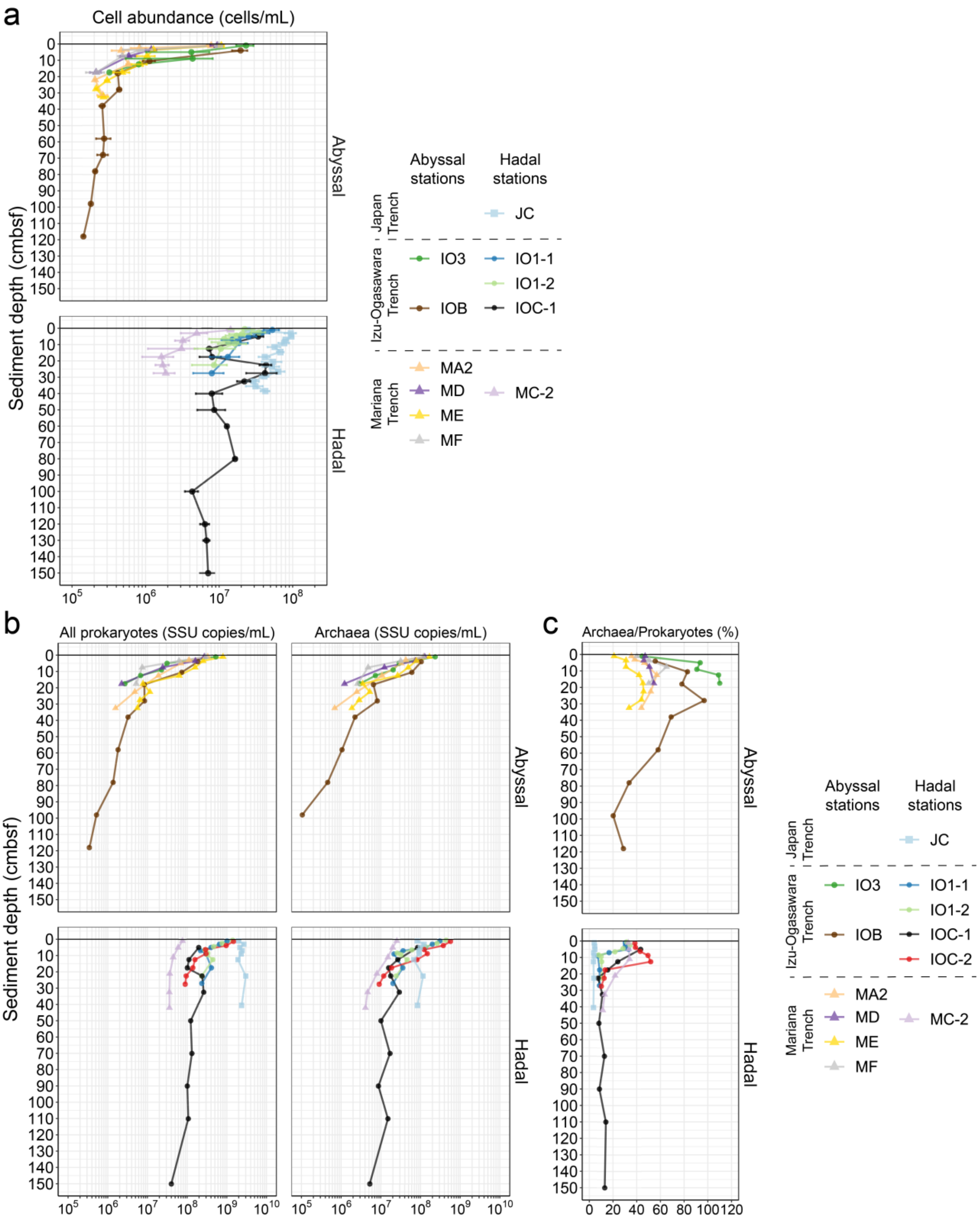


**Figure S4.** The abundance of microbes and ratios of archaea in each sediment core from abyssal (upper panels) and hadal (lower panels) stations. The abundances were measured using (**a**) cell counting and (**b**) qPCR techniques. The X-axes represent cell counts and SSU rRNA gene copies per milliliter of sediment, respectively. The error bars represent standard deviation. (**c**) Ratios of archaea/prokaryotes were calculated using qPCR data.

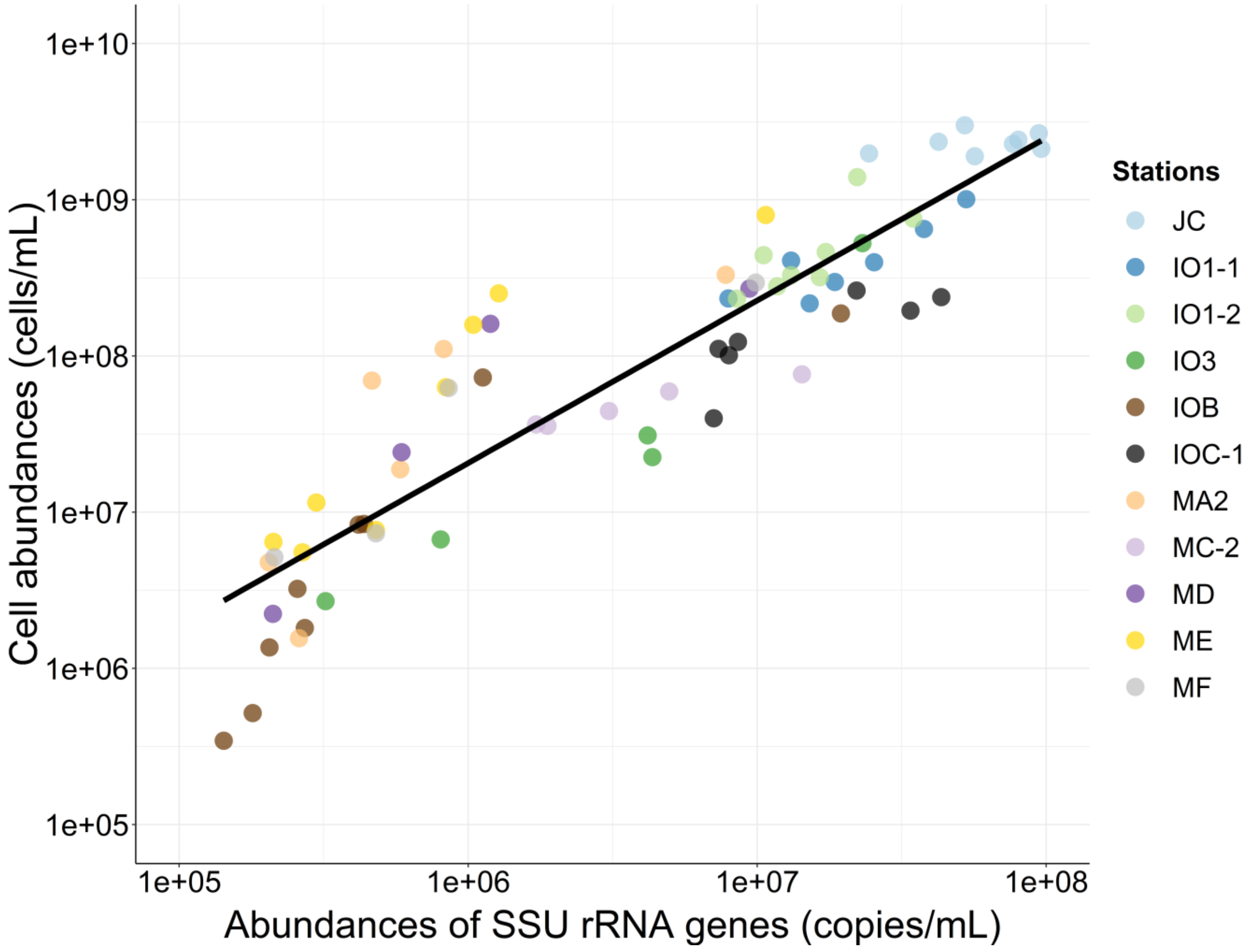


**Figure S5.** Relationship between qPCR counts and cell counts. Only sediment samples in which both cell counts and qPCR were conducted were included in this analysis. The equation of the linear regression line was y = 28.2x + 4.6e+06 ($r^2$=0.75). Each sediment core is coded by color.

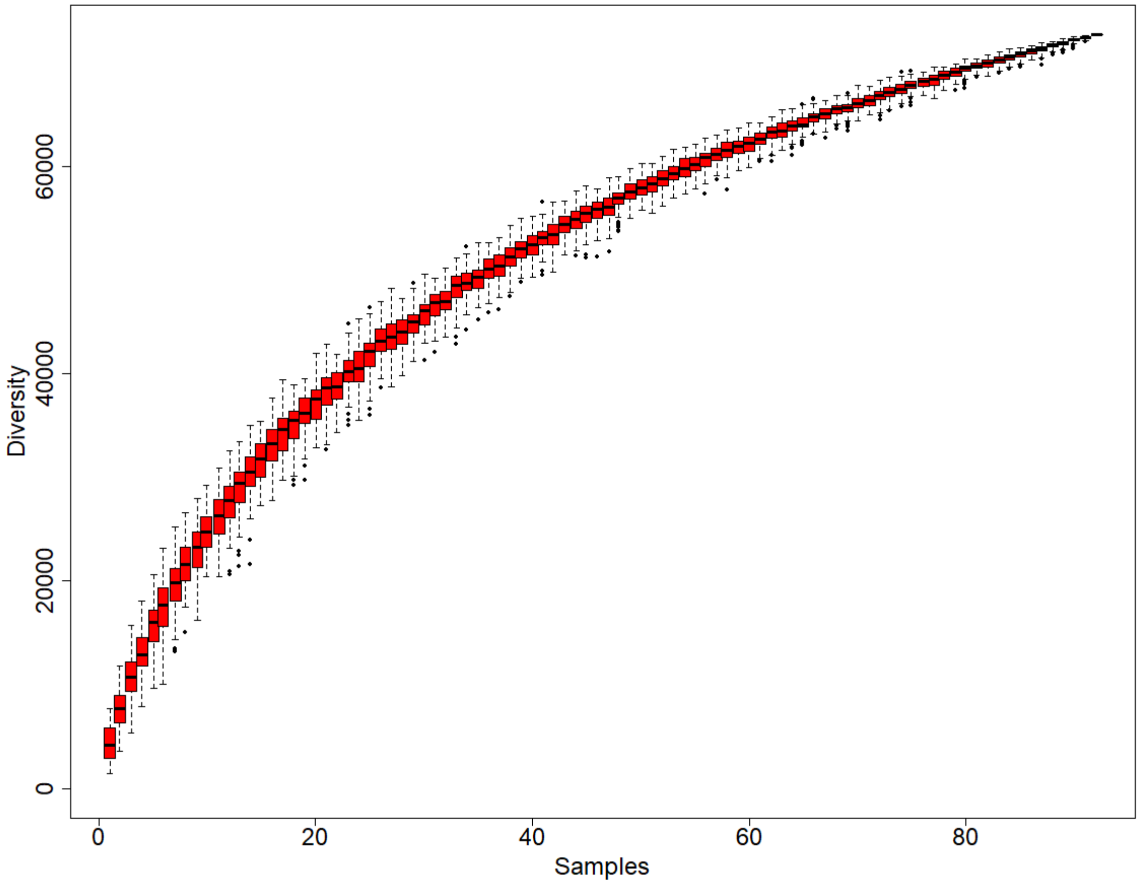


**Figure S6.** Rarefaction curves of the OTU numbers (diversity) against the number of examined samples.

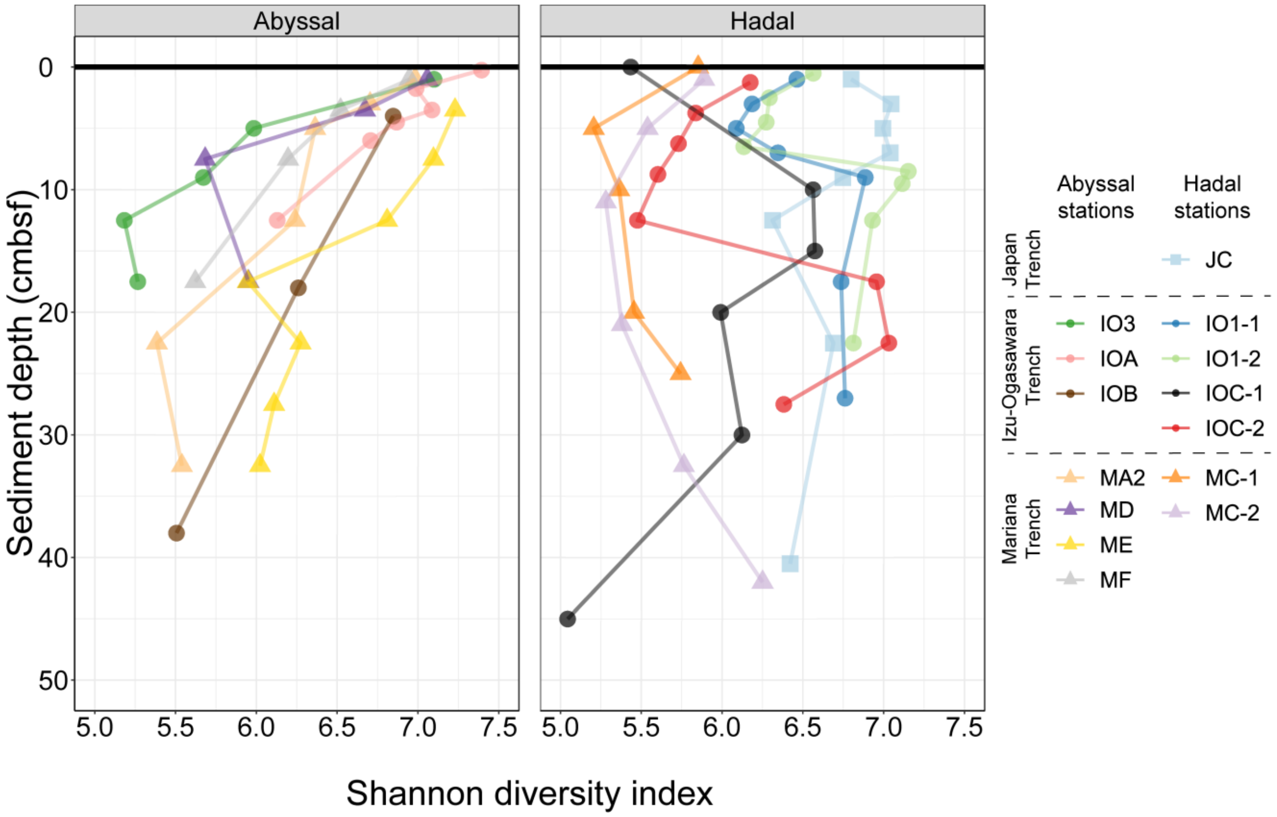


**Figure S7.** Shannon diversity index at the OTU level versus sediment depth from abyssal (left panel) and hadal (right panel) stations. The colored lines represent the values of each sampling station. Data from layers ranged between 0 to 50 cmbsf are shown in this figure.

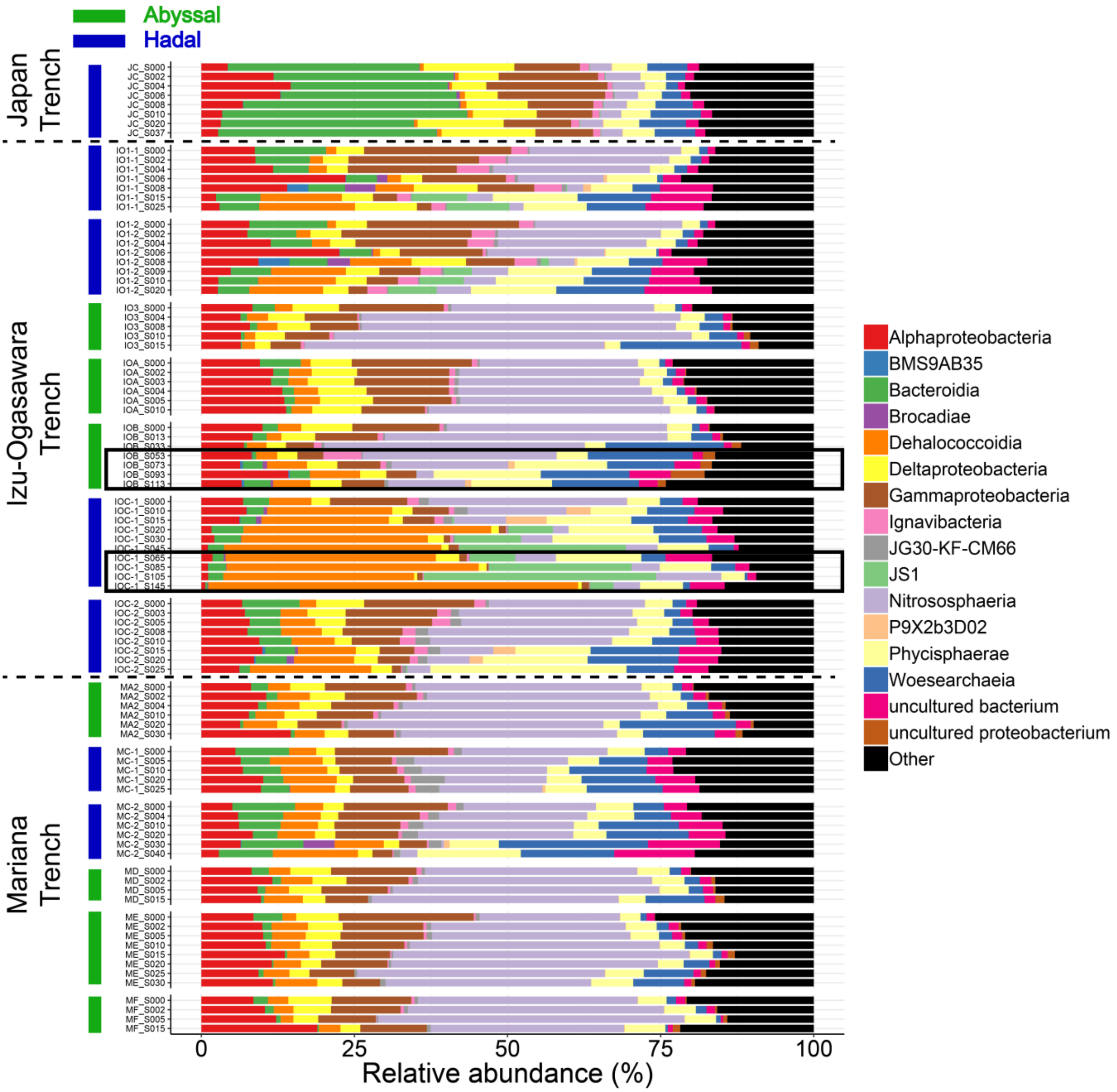


**Figure S8.** Relative abundances of sequences at the class level. Groups demonstrating <5% abundance are summarized as 'Other'. Sediment samples retrieved from deep sediment layers (>s50 cmbsf) are indicated by surrounding black rectangles.

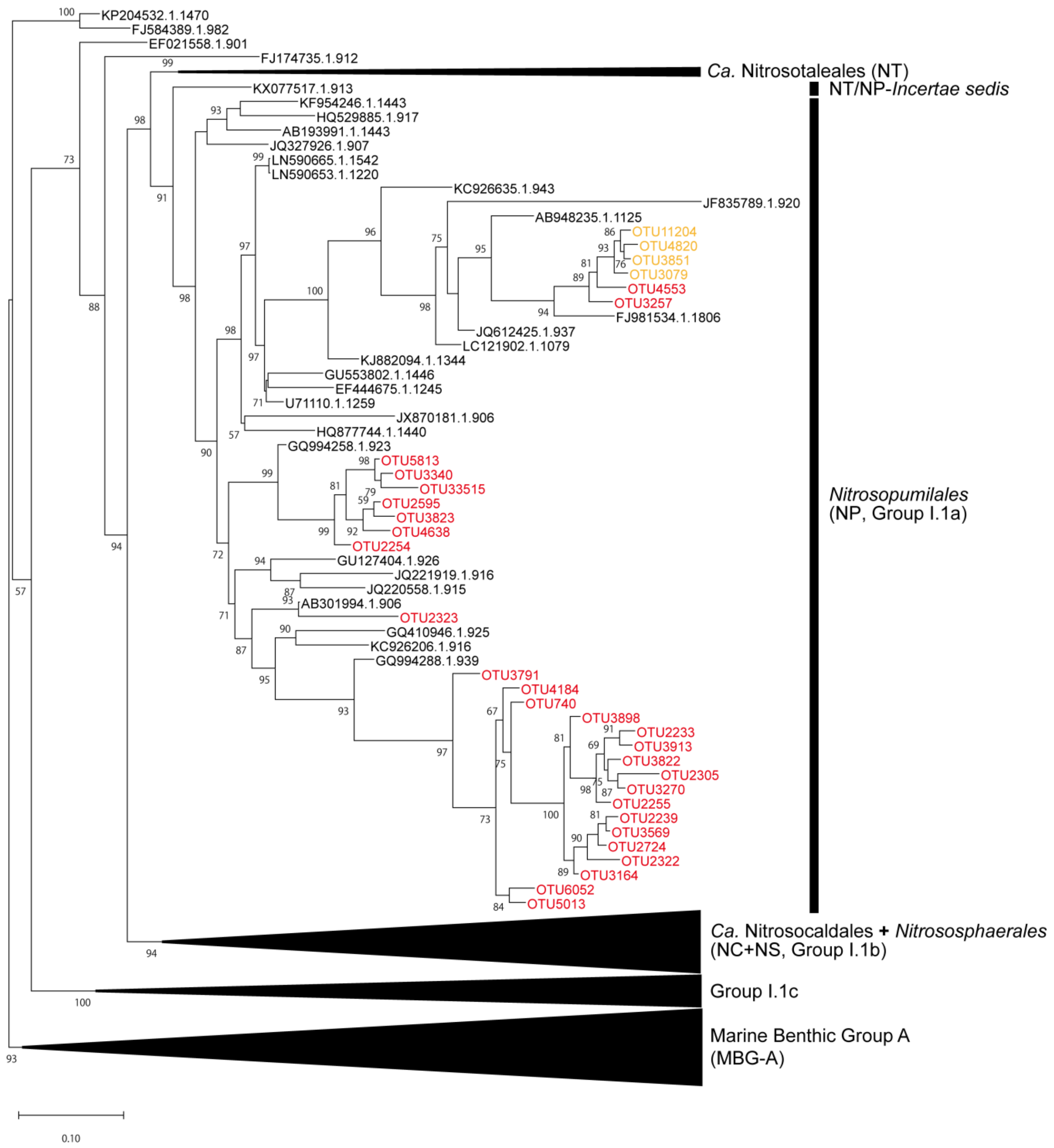


**Figure S9.** Phylogenetic tree of Thaumarchaeota SSU rRNA gene sequences. Nodes colored by red and orange represent OTUs belonging to the co-occurrence group A and D, respectively. Numbers adjusted with edges indicate bootstrap supports, and only values >50 are shown. Scale bar represents the estimated number of substitutions per site.

### Supplementary Tables

**Table S1.** Station descriptions and sequencing statistics of each sediment sample

| Sample | Cruise ID | Collection date | Sampling method | Area | Description | Site | Sation | Location | Environmental feature | Elevation (mbsl) | Depth range (cmbsf) | Sequenced reads | OTUs | Sequencing data accession | Reference |
|---|---|---|---|---|---|---|---|---|---|---|---|---|---|---|---|
| JC_S000 | KR12-19 | 2012-12-03 | 11K Lander system | Japan Trench | JC | J36N | J36N | 36.0724 N 142.7440 E | Hadalpelagic trench bottom | 7,963 | 0–2 | 63,396 | 7,508 | SAMD00165472 | - |
| JC_S002 | KR12-19 | 2012-12-03 | 11K Lander system | Japan Trench | JC | J36N | J36N | 36.0724 N 142.7440 E | Hadalpelagic trench bottom | 7,963 | 2–4 | 63,647 | 7,026 | SAMD00165473 | - |
| JC_S004 | KR12-19 | 2012-12-03 | 11K Lander system | Japan Trench | JC | J36N | J36N | 36.0724 N 142.7440 E | Hadalpelagic trench bottom | 7,963 | 4–6 | 62,583 | 6,507 | SAMD00165474 | - |
| JC_S006 | KR12-19 | 2012-12-03 | 11K Lander system | Japan Trench | JC | J36N | J36N | 36.0724 N 142.7440 E | Hadalpelagic trench bottom | 7,963 | 6–8 | 69,529 | 7,089 | SAMD00165475 | - |
| JC_S008 | KR12-19 | 2012-12-03 | 11K Lander system | Japan Trench | JC | J36N | J36N | 36.0724 N 142.7440 E | Hadalpelagic trench bottom | 7,963 | 8–10 | 64,605 | 7,184 | SAMD00165476 | - |
| JC_S010 | KR12-19 | 2012-12-03 | 11K Lander system | Japan Trench | JC | J36N | J36N | 36.0724 N 142.7440 E | Hadalpelagic trench bottom | 7,963 | 10–15 | 53,895 | 5,934 | SAMD00165477 | - |
| JC_S020 | KR12-19 | 2012-12-03 | 11K Lander system | Japan Trench | JC | J36N | J36N | 36.0724 N 142.7440 E | Hadalpelagic trench bottom | 7,963 | 20–25 | 65,720 | 7,225 | SAMD00165478 | - |
| JC_S037 | KR12-19 | 2012-12-03 | 11K Lander system | Japan Trench | JC | J36N | J36N | 36.0724 N 142.7440 E | Hadalpelagic trench bottom | 7,963 | 37–44 | 65,526 | 6,870 | SAMD00165479 | - |
| IO1-1_S000 | KR15-01 | 2015-03-15 | 11K Lander system | Izu Ogasawara Trench | IO1-1 | IO1 | IO1-1 | 33.7500 N 142.0045 E | Hadalpelagic trench bottom | 9,250 | 0–2 | 81,741 | 6,055 | SAMD00165421 | - |
| IO1-1_S002 | KR15-01 | 2015-03-15 | 11K Lander system | Izu Ogasawara Trench | IO1-1 | IO1 | IO1-1 | 33.7500 N 142.0045 E | Hadalpelagic trench bottom | 9,250 | 2–4 | 82,347 | 5,675 | SAMD00165422 | - |
| IO1-1_S004 | KR15-01 | 2015-03-15 | 11K Lander system | Izu Ogasawara Trench | IO1-1 | IO1 | IO1-1 | 33.7500 N 142.0045 E | Hadalpelagic trench bottom | 9,250 | 4–6 | 76,500 | 5,178 | SAMD00165423 | - |
| IO1-1_S006 | KR15-01 | 2015-03-15 | 11K Lander system | Izu Ogasawara Trench | IO1-1 | IO1 | IO1-1 | 33.7500 N 142.0045 E | Hadalpelagic trench bottom | 9,250 | 6–8 | 74,935 | 4,278 | SAMD00165424 | - |
| IO1-1_S008 | KR15-01 | 2015-03-15 | 11K Lander system | Izu Ogasawara Trench | IO1-1 | IO1 | IO1-1 | 33.7500 N 142.0045 E | Hadalpelagic trench bottom | 9,250 | 8–10 | 72,613 | 6,457 | SAMD00165425 | - |
| IO1-1_S015 | KR15-01 | 2015-03-15 | 11K Lander system | Izu Ogasawara Trench | IO1-1 | IO1 | IO1-1 | 33.7500 N 142.0045 E | Hadalpelagic trench bottom | 9,250 | 15–20 | 57,460 | 6,231 | SAMD00165426 | - |
| IO1-1_S025 | KR15-01 | 2015-03-15 | 11K Lander system | Izu Ogasawara Trench | IO1-1 | IO1 | IO1-1 | 33.7500 N 142.0045 E | Hadalpelagic trench bottom | 9,250 | 25–29 | 70,130 | 6,096 | SAMD00165427 | - |
| IO1-2_S000 | KR15-01 | 2015-03-18 | 11K Lander system | Izu Ogasawara Trench | IO1-2 | IO1 | IO1-2 | 33.7488 N 142.0003 E | Hadalpelagic trench bottom | 9,242 | 0–1 | 77,885 | 6,466 | SAMD00165428 | - |
| IO1-2_S002 | KR15-01 | 2015-03-18 | 11K Lander system | Izu Ogasawara Trench | IO1-2 | IO1 | IO1-2 | 33.7488 N 142.0003 E | Hadalpelagic trench bottom | 9,242 | 2–3 | 80,540 | 5,850 | SAMD00165429 | - |
| IO1-2_S004 | KR15-01 | 2015-03-18 | 11K Lander system | Izu Ogasawara Trench | IO1-2 | IO1 | IO1-2 | 33.7488 N 142.0003 E | Hadalpelagic trench bottom | 9,242 | 4–5 | 76,858 | 5,566 | SAMD00165430 | - |
| IO1-2_S006 | KR15-01 | 2015-03-18 | 11K Lander system | Izu Ogasawara Trench | IO1-2 | IO1 | IO1-2 | 33.7488 N 142.0003 E | Hadalpelagic trench bottom | 9,242 | 6–7 | 87,265 | 4,007 | SAMD00165431 | - |
| IO1-2_S008 | KR15-01 | 2015-03-18 | 11K Lander system | Izu Ogasawara Trench | IO1-2 | IO1 | IO1-2 | 33.7488 N 142.0003 E | Hadalpelagic trench bottom | 9,242 | 8–9 | 75,608 | 7,318 | SAMD00165432 | - |
| IO1-2_S009 | KR15-01 | 2015-03-18 | 11K Lander system | Izu Ogasawara Trench | IO1-2 | IO1 | IO1-2 | 33.7488 N 142.0003 E | Hadalpelagic trench bottom | 9,242 | 9–10 | 73,339 | 7,206 | SAMD00165433 | - |
| IO1-2_S010 | KR15-01 | 2015-03-18 | 11K Lander system | Izu Ogasawara Trench | IO1-2 | IO1 | IO1-2 | 33.7488 N 142.0003 E | Hadalpelagic trench bottom | 9,242 | 10–15 | 72,303 | 7,094 | SAMD00165434 | - |
| IO1-2_S020 | KR15-01 | 2015-03-18 | 11K Lander system | Izu Ogasawara Trench | IO1-2 | IO1 | IO1-2 | 33.7488 N 142.0003 E | Hadalpelagic trench bottom | 9,242 | 20–25 | 65,868 | 6,124 | SAMD00165435 | - |
| IO3_S000 | KR15-01 | 2015-03-16 | 11K Lander system | Izu Ogasawara Trench | IO3 | IO3 | IO3 | 33.7434 N 142.8830 E | Abyssalpelagic plain | 5,257 | 0–2 | 74,654 | 8,723 | SAMD00165436 | - |
| IO3_S004 | KR15-01 | 2015-03-16 | 11K Lander system | Izu Ogasawara Trench | IO3 | IO3 | IO3 | 33.7434 N 142.8830 E | Abyssalpelagic plain | 5,257 | 4–6 | 81,956 | 4,917 | SAMD00165437 | - |
| IO3_S008 | KR15-01 | 2015-03-16 | 11K Lander system | Izu Ogasawara Trench | IO3 | IO3 | IO3 | 33.7434 N 142.8830 E | Abyssalpelagic plain | 5,257 | 8–10 | 75,728 | 3,271 | SAMD00165438 | - |
| IO3_S010 | KR15-01 | 2015-03-16 | 11K Lander system | Izu Ogasawara Trench | IO3 | IO3 | IO3 | 33.7434 N 142.8830 E | Abyssalpelagic plain | 5,257 | 10–15 | 78,988 | 2,543 | SAMD00165439 | - |
| IO3_S015 | KR15-01 | 2015-03-16 | 11K Lander system | Izu Ogasawara Trench | IO3 | IO3 | IO3 | 33.7434 N 142.8830 E | Abyssalpelagic plain | 5,257 | 15–20 | 69,268 | 2,034 | SAMD00165440 | - |
| IOA_S000 | YK11-06 | 2011-09-07 | Shinkai6500 & push corer | Izu Ogasawara Trench | IOA | 30N | 30N | 30.1508 N 143.5832 E | Abyssalpelagic plain | 5,370 | 0–0.5 | 105,234 | 10,703 | SAMD00165441 | - |
| IOA_S002 | YK11-06 | 2011-09-07 | Shinkai6500 & push corer | Izu Ogasawara Trench | IOA | 30N | 30N | 30.1508 N 143.5832 E | Abyssalpelagic plain | 5,370 | 1.5–2 | 101,817 | 9,276 | SAMD00165442 | - |
| IOA_S003 | YK11-06 | 2011-09-07 | Shinkai6500 & push corer | Izu Ogasawara Trench | IOA | 30N | 30N | 30.1508 N 143.5832 E | Abyssalpelagic plain | 5,370 | 3–4 | 103,968 | 10,252 | SAMD00165443 | - |
| IOA_S004 | YK11-06 | 2011-09-07 | Shinkai6500 & push corer | Izu Ogasawara Trench | IOA | 30N | 30N | 30.1508 N 143.5832 E | Abyssalpelagic plain | 5,370 | 4–5 | 120,851 | 9,673 | SAMD00165444 | - |
| IOA_S005 | YK11-06 | 2011-09-07 | Shinkai6500 & push corer | Izu Ogasawara Trench | IOA | 30N | 30N | 30.1508 N 143.5832 E | Abyssalpelagic plain | 5,370 | 5–7 | 111,661 | 8,991 | SAMD00165445 | - |
| IOA_S010 | YK11-06 | 2011-09-07 | Shinkai6500 & push corer | Izu Ogasawara Trench | IOA | 30N | 30N | 30.1508 N 143.5832 E | Abyssalpelagic plain | 5,370 | 10–15 | 130,129 | 5,645 | SAMD00165446 | - |
| IOB_S000 | KR11-11 | 2011-12-12 | ABISMO & gravity corer | Izu Ogasawara Trench | IOB | IOAB16 | IOAB16 | 29.2746 N 143.7673 E | Abyssalpelagic plain | 5,474 | 0–8 | 74,868 | 8,989 | SAMD00165447 | - |
| IOB_S013 | KR11-11 | 2011-12-12 | ABISMO & gravity corer | Izu Ogasawara Trench | IOB | IOAB16 | IOAB16 | 29.2746 N 143.7673 E | Abyssalpelagic plain | 5,474 | 13–23 | 78,714 | 5,189 | SAMD00165448 | - |
| IOB_S033 | KR11-11 | 2011-12-12 | ABISMO & gravity corer | Izu Ogasawara Trench | IOB | IOAB16 | IOAB16 | 29.2746 N 143.7673 E | Abyssalpelagic plain | 5,474 | 33–43 | 70,566 | 2,935 | SAMD00165449 | - |
| IOB_S053 | KR11-11 | 2011-12-12 | ABISMO & gravity corer | Izu Ogasawara Trench | IOB | IOAB16 | IOAB16 | 29.2746 N 143.7673 E | Abyssalpelagic plain | 5,474 | 53–63 | 74,210 | 2,617 | SAMD00165450 | - |
| IOB_S073 | KR11-11 | 2011-12-12 | ABISMO & gravity corer | Izu Ogasawara Trench | IOB | IOAB16 | IOAB16 | 29.2746 N 143.7673 E | Abyssalpelagic plain | 5,474 | 73–83 | 74,301 | 3,461 | SAMD00165451 | - |
| IOB_S093 | KR11-11 | 2011-12-12 | ABISMO & gravity corer | Izu Ogasawara Trench | IOB | IOAB16 | IOAB16 | 29.2746 N 143.7673 E | Abyssalpelagic plain | 5,474 | 93–103 | 61,038 | 1,587 | SAMD00165452 | - |
| IOB_S113 | KR11-11 | 2011-12-12 | ABISMO & gravity corer | Izu Ogasawara Trench | IOB | IOAB16 | IOAB16 | 29.2746 N 143.7673 E | Abyssalpelagic plain | 5,474 | 113–123 | 68,945 | 1,806 | SAMD00165453 | - |
| IOC-1_S000 | KR11-11 | 2011-12-11 | ABISMO & gravity corer | Izu Ogasawara Trench | IOC-1 | IOAB15 | IOAB15-1 | 29.1500 N 142.8033 E | Hadalpelagic trench bottom | 9,776 | 0–10 | 118,398 | 3,413 | SAMD00165454 | - |
| IOC-1_S010 | KR11-11 | 2011-12-11 | ABISMO & gravity corer | Izu Ogasawara Trench | IOC-1 | IOAB15 | IOAB15-1 | 29.1500 N 142.8033 E | Hadalpelagic trench bottom | 9,776 | 10–15 | 111,724 | 3,959 | SAMD00165455 | - |
| IOC-1_S015 | KR11-11 | 2011-12-11 | ABISMO & gravity corer | Izu Ogasawara Trench | IOC-1 | IOAB15 | IOAB15-1 | 29.1500 N 142.8033 E | Hadalpelagic trench bottom | 9,776 | 15–20 | 116,545 | 3,825 | SAMD00165456 | - |
| IOC-1_S020 | KR11-11 | 2011-12-11 | ABISMO & gravity corer | Izu Ogasawara Trench | IOC-1 | IOAB15 | IOAB15-1 | 29.1500 N 142.8033 E | Hadalpelagic trench bottom | 9,776 | 20–25 | 134,679 | 3,619 | SAMD00165457 | - |
| IOC-1_S030 | KR11-11 | 2011-12-11 | ABISMO & gravity corer | Izu Ogasawara Trench | IOC-1 | IOAB15 | IOAB15-1 | 29.1500 N 142.8033 E | Hadalpelagic trench bottom | 9,776 | 30–35 | 121,166 | 4,053 | SAMD00165458 | - |
| IOC-1_S045 | KR11-11 | 2011-12-11 | ABISMO & gravity corer | Izu Ogasawara Trench | IOC-1 | IOAB15 | IOAB15-1 | 29.1500 N 142.8033 E | Hadalpelagic trench bottom | 9,776 | 45–55 | 113,299 | 3,060 | SAMD00165459 | - |
| IOC-1_S065 | KR11-11 | 2011-12-11 | ABISMO & gravity corer | Izu Ogasawara Trench | IOC-1 | IOAB15 | IOAB15-1 | 29.1500 N 142.8033 E | Hadalpelagic trench bottom | 9,776 | 65–75 | 124,060 | 3,972 | SAMD00165460 | - |
| IOC-1_S085 | KR11-11 | 2011-12-11 | ABISMO & gravity corer | Izu Ogasawara Trench | IOC-1 | IOAB15 | IOAB15-1 | 29.1500 N 142.8033 E | Hadalpelagic trench bottom | 9,776 | 85–95 | 114,250 | 2,967 | SAMD00165461 | - |
| IOC-1_S105 | KR11-11 | 2011-12-11 | ABISMO & gravity corer | Izu Ogasawara Trench | IOC-1 | IOAB15 | IOAB15-1 | 29.1500 N 142.8033 E | Hadalpelagic trench bottom | 9,776 | 105–115 | 124,747 | 2,600 | SAMD00165462 | - |
| IOC-1_S145 | KR11-11 | 2011-12-11 | ABISMO & gravity corer | Izu Ogasawara Trench | IOC-1 | IOAB15 | IOAB15-1 | 29.1500 N 142.8033 E | Hadalpelagic trench bottom | 9,776 | 145–155 | 117,905 | 2,386 | SAMD00165463 | - |
| IOC-2_S000 | KR11-11 | 2011-12-23 | 11K Lander system | Izu Ogasawara Trench | IOC-2 | IOAB15 | IOAB15-2 | 29.1412 N 142.7983 E | Hadalpelagic trench bottom | 9,772 | 0–2.5 | 74,684 | 5,852 | SAMD00165464 | - |
| IOC-2_S003 | KR11-11 | 2011-12-23 | 11K Lander system | Izu Ogasawara Trench | IOC-2 | IOAB15 | IOAB15-2 | 29.1412 N 142.7983 E | Hadalpelagic trench bottom | 9,772 | 2.5–5 | 64,424 | 4,760 | SAMD00165465 | - |
| IOC-2_S005 | KR11-11 | 2011-12-23 | 11K Lander system | Izu Ogasawara Trench | IOC-2 | IOAB15 | IOAB15-2 | 29.1412 N 142.7983 E | Hadalpelagic trench bottom | 9,772 | 5–7.5 | 65,632 | 4,085 | SAMD00165466 | - |
| IOC-2_S008 | KR11-11 | 2011-12-23 | 11K Lander system | Izu Ogasawara Trench | IOC-2 | IOAB15 | IOAB15-2 | 29.1412 N 142.7983 E | Hadalpelagic trench bottom | 9,772 | 7.5–10 | 76,150 | 4,839 | SAMD00165467 | - |
| IOC-2_S010 | KR11-11 | 2011-12-23 | 11K Lander system | Izu Ogasawara Trench | IOC-2 | IOAB15 | IOAB15-2 | 29.1412 N 142.7983 E | Hadalpelagic trench bottom | 9,772 | 10–15 | 77,633 | 4,479 | SAMD00165468 | - |
| IOC-2_S015 | KR11-11 | 2011-12-23 | 11K Lander system | Izu Ogasawara Trench | IOC-2 | IOAB15 | IOAB15-2 | 29.1412 N 142.7983 E | Hadalpelagic trench bottom | 9,772 | 15–20 | 52,922 | 5,341 | SAMD00165469 | - |
| IOC-2_S020 | KR11-11 | 2011-12-23 | 11K Lander system | Izu Ogasawara Trench | IOC-2 | IOAB15 | IOAB15-2 | 29.1412 N 142.7983 E | Hadalpelagic trench bottom | 9,772 | 20–25 | 56,469 | 5,483 | SAMD00165470 | - |
| IOC-2_S025 | KR11-11 | 2011-12-23 | 11K Lander system | Izu Ogasawara Trench | IOC-2 | IOAB15 | IOAB15-2 | 29.1412 N 142.7983 E | Hadalpelagic trench bottom | 9,772 | 25–30 | 54,921 | 4,092 | SAMD00165471 | - |
| MA2_S000 | KR14-01 | 2014-01-18 | Multiple corer | Mariana Trench | MA2 | MA2 | MA2 | 11.7463 N 142.1087 E | Abyssalpelagic trench slope | 5,838 | 0–2 | 65,551 | 8,549 | SAMD00078379 | Hirai et al. 2018 |
| MA2_S002 | KR14-01 | 2014-01-18 | Multiple corer | Mariana Trench | MA2 | MA2 | MA2 | 11.7463 N 142.1087 E | Abyssalpelagic trench slope | 5,838 | 2–4 | 63,157 | 6,596 | SAMD00165480 | - |
| MA2_S004 | KR14-01 | 2014-01-18 | Multiple corer | Mariana Trench | MA2 | MA2 | MA2 | 11.7463 N 142.1087 E | Abyssalpelagic trench slope | 5,838 | 4–6 | 60,170 | 5,459 | SAMD00165481 | - |
| MA2_S010 | KR14-01 | 2014-01-18 | Multiple corer | Mariana Trench | MA2 | MA2 | MA2 | 11.7463 N 142.1087 E | Abyssalpelagic trench slope | 5,838 | 10–15 | 68,203 | 6,064 | SAMD00165482 | - |
| MA2_S020 | KR14-01 | 2014-01-18 | Multiple corer | Mariana Trench | MA2 | MA2 | MA2 | 11.7463 N 142.1087 E | Abyssalpelagic trench slope | 5,838 | 20–25 | 69,698 | 3,503 | SAMD00165483 | - |
| MA2_S030 | KR14-01 | 2014-01-18 | Multiple corer | Mariana Trench | MA2 | MA2 | MA2 | 11.7463 N 142.1087 E | Abyssalpelagic trench slope | 5,838 | 30–35 | 65,332 | 3,268 | SAMD00165484 | - |
| MC-1_S000 | KR14-01 | 2014-01-12 | 11K Lander system | Mariana Trench | MC-1 | MC | MC-1 | 11.3721 N 142.4378 E | Hadalpelagic trench bottom | 10,901 | 0–2 | 125,322 | 4,286 | SAMD00165485 | - |
| MC-1_S005 | KR14-01 | 2014-01-12 | 11K Lander system | Mariana Trench | MC-1 | MC | MC-1 | 11.3721 N 142.4378 E | Hadalpelagic trench bottom | 10,901 | 5–10 | 116,070 | 2,796 | SAMD00165486 | - |
| MC-1_S010 | KR14-01 | 2014-01-12 | 11K Lander system | Mariana Trench | MC-1 | MC | MC-1 | 11.3721 N 142.4378 E | Hadalpelagic trench bottom | 10,901 | 10–15 | 108,498 | 2,768 | SAMD00165487 | - |
| MC-1_S020 | KR14-01 | 2014-01-12 | 11K Lander system | Mariana Trench | MC-1 | MC | MC-1 | 11.3721 N 142.4378 E | Hadalpelagic trench bottom | 10,901 | 20–25 | 121,610 | 2,791 | SAMD00165488 | - |
| MC-1_S025 | KR14-01 | 2014-01-12 | 11K Lander system | Mariana Trench | MC-1 | MC | MC-1 | 11.3721 N 142.4378 E | Hadalpelagic trench bottom | 10,901 | 25–30 | 119,924 | 3,777 | SAMD00165489 | - |
| MC-2_S000 | KR14-01 | 2014-01-17 | 11K Lander system | Mariana Trench | MC-2 | MC | MC-2 | 11.3699 N 142.4293 E | Hadalpelagic trench bottom | 10,902 | 0–2 | 69,377 | 4,037 | SAMD00165490 | - |
| MC-2_S004 | KR14-01 | 2014-01-17 | 11K Lander system | Mariana Trench | MC-2 | MC | MC-2 | 11.3699 N 142.4293 E | Hadalpelagic trench bottom | 10,902 | 4–6 | 68,850 | 3,571 | SAMD00165491 | - |
| MC-2_S010 | KR14-01 | 2014-01-17 | 11K Lander system | Mariana Trench | MC-2 | MC | MC-2 | 11.3699 N 142.4293 E | Hadalpelagic trench bottom | 10,902 | 10–12 | 63,596 | 3,119 | SAMD00165492 | - |
| MC-2_S020 | KR14-01 | 2014-01-17 | 11K Lander system | Mariana Trench | MC-2 | MC | MC-2 | 11.3699 N 142.4293 E | Hadalpelagic trench bottom | 10,902 | 20–22 | 67,648 | 3,655 | SAMD00165493 | - |
| MC-2_S030 | KR14-01 | 2014-01-17 | 11K Lander system | Mariana Trench | MC-2 | MC | MC-2 | 11.3699 N 142.4293 E | Hadalpelagic trench bottom | 10,902 | 30–35 | 67,786 | 3,601 | SAMD00165494 | - |
| MC-2_S040 | KR14-01 | 2014-01-17 | 11K Lander system | Mariana Trench | MC-2 | MC | MC-2 | 11.3699 N 142.4293 E | Hadalpelagic trench bottom | 10,902 | 40–44 | 44,215 | 2,985 | SAMD00165495 | - |
| MD_S000 | KR14-01 | 2014-01-13 | Multiple corer | Mariana Trench | MD | MD | MD | 10.9158 N 142.5339 E | Abyssalpelagic trench slope | 6,067 | 0–2 | 69,332 | 8,520 | SAMD00169212 | - |
| MD_S002 | KR14-01 | 2014-01-13 | Multiple corer | Mariana Trench | MD | MD | MD | 10.9158 N 142.5339 E | Abyssalpelagic trench slope | 6,067 | 2–5 | 61,410 | 6,310 | SAMD00169213 | - |
| MD_S005 | KR14-01 | 2014-01-13 | Multiple corer | Mariana Trench | MD | MD | MD | 10.9158 N 142.5339 E | Abyssalpelagic trench slope | 6,067 | 5–10 | 67,597 | 3,489 | SAMD00169214 | - |
| MD_S015 | KR14-01 | 2014-01-13 | Multiple corer | Mariana Trench | MD | MD | MD | 10.9158 N 142.5339 E | Abyssalpelagic trench slope | 6,067 | 15–20 | 77,711 | 3,548 | SAMD00169215 | - |
| ME_S000 | KR14-01 | 2014-01-13 | Multiple corer | Mariana Trench | ME | ME | ME | 10.2997 N 142.6003E | Abyssalpelagic plain | 4,700 | 0–2 | 196,691 | 13,181 | SAMD00165500 | - |
| ME_S002 | KR14-01 | 2014-01-13 | Multiple corer | Mariana Trench | ME | ME | ME | 10.2997 N 142.6003 E | Abyssalpelagic plain | 4,700 | 5–10 | 191,271 | 12,406 | SAMD00165501 | - |
| ME_S005 | KR14-01 | 2014-01-13 | Multiple corer | Mariana Trench | ME | ME | ME | 10.2997 N 142.6003 E | Abyssalpelagic plain | 4,700 | 10–15 | 177,753 | 10,530 | SAMD00165502 | - |
| ME_S010 | KR14-01 | 2014-01-13 | Multiple corer | Mariana Trench | ME | ME | ME | 10.2997 N 142.6003 E | Abyssalpelagic plain | 4,700 | 15–20 | 154,996 | 9,087 | SAMD00165503 | - |
| ME_S015 | KR14-01 | 2014-01-13 | Multiple corer | Mariana Trench | ME | ME | ME | 10.2997 N 142.6003 E | Abyssalpelagic plain | 4,700 | 20–25 | 215,201 | 6,183 | SAMD00165504 | - |
| ME_S020 | KR14-01 | 2014-01-13 | Multiple corer | Mariana Trench | ME | ME | ME | 10.2997 N 142.6003 E | Abyssalpelagic plain | 4,700 | 25–30 | 178,005 | 6,617 | SAMD00165505 | - |
| ME_S025 | KR14-01 | 2014-01-13 | Multiple corer | Mariana Trench | ME | ME | ME | 10.2997 N 142.6003 E | Abyssalpelagic plain | 4,700 | 30–35 | 157,052 | 5,013 | SAMD00165506 | - |
| ME_S030 | KR14-01 | 2014-01-13 | Multiple corer | Mariana Trench | ME | ME | ME | 10.2997 N 142.6003 E | Abyssalpelagic plain | 4,700 | 2–5 | 184,610 | 4,725 | SAMD00165507 | - |
| MF_S000 | KR14-01 | 2014-01-14 | Multiple corer | Mariana Trench | MF | MF | MF | 10.6499 N 142.5512 E | Abyssalpelagic plain | 5,183 | 0–2 | 67,186 | 8,376 | SAMD00169216 | - |
| MF_S002 | KR14-01 | 2014-01-14 | Multiple corer | Mariana Trench | MF | MF | MF | 10.6499 N 142.5512 E | Abyssalpelagic plain | 5,183 | 2–5 | 68,994 | 6,885 | SAMD00169217 | - |
| MF_S005 | KR14-01 | 2014-01-14 | Multiple corer | Mariana Trench | MF | MF | MF | 10.6499 N 142.5512 E | Abyssalpelagic plain | 5,183 | 5–10 | 71,778 | 4,612 | SAMD00169218 | - |
| MF_S015 | KR14-01 | 2014-01-14 | Multiple corer | Mariana Trench | MF | MF | MF | 10.6499 N 142.5512 E | Abyssalpelagic plain | 5,183 | 15–20 | 71,117 | 3,089 | SAMD00169219 | - |

**Table S2.** Primers, probes, and amplification conditions of qPCR analyses

| Target gene | Primer name | Primer sequence (5'–3') | Probe name | Probe sequence (5'–3') | PCR condition |
|---|---|---|---|---|---|
| Prokaryotic SSU rRNA gene | Uni340F | CCTACGGGRBGCASCAG | Uni516F | TGYCAGCMGCCGCGGTAAHACVNRS | 96°C for 1min, 50×(96°C 25s, 57°C 6 min) (qPCR Quick GoldStar Mastermix Plus) |
| | Uni806R | GGACTACNNGGGTATCTAAT | | | 96°C for 1min, 50×(96°C 25s, 57°C 2 min) (Premix Ex Taq [Perfect Real Time]) |
| Archaeal SSU rRNA gene | Arch349F | GYGCASCAGKCGMGAAW | Arch516F | TGYCAGCCGCCGCGGTAAHACCVGC | 96°C for 1min, 50×(96°C 25s, 59°C 6 min) (qPCR Quick GoldStar Mastermix Plus) |
| | Arch806R | GGACTACVSGGGTATCTAAT | | | 96°C for 1min, 50×(96°C 25s, 58°C 2 min) (Premix Ex Taq [Perfect Real Time]) |

F: forward primer; R: reverse primer

**Table S3.** Primers and adapters used for SSU rRNA gene PCR amplification

| Primer name | P7/P5/Seq adapter | Primer sequence (5'–3') | Mix rate |
|---|---|---|---|
| Uni530F1N_TruSecP5side | ACACTCTTTCCCTACACGACGCTCTTCCGATCT | GTGCCAGCAGCCGCGG | 30 |
| Uni530F2N_TruSecP5side | ACACTCTTTCCCTACACGACGCTCTTCCGATCT | GTGBCAGCCGCCGCGG | 24 |
| Uni530F3N_TruSecP5side | ACACTCTTTCCCTACACGACGCTCTTCCGATCT | YTGCCAGCCGCCGCGG | 6 |
| Uni530F4N_TruSecP5side | ACACTCTTTCCCTACACGACGCTCTTCCGATCT | GTGCCAGCAGCWGCGG | 1 |
| Uni530F5N_TruSecP5side | ACACTCTTTCCCTACACGACGCTCTTCCGATCT | GTGCCAGCAGTCGCGG | 1 |
| Uni530F6N_TruSecP5side | ACACTCTTTCCCTACACGACGCTCTTCCGATCT | GTGCCAGAAGMMTCGG | 1 |
| Uni530F7N_TruSecP5side | ACACTCTTTCCCTACACGACGCTCTTCCGATCT | GTGGCAGTCGCCACGG | 3 |
| Uni907R1N_TruSecP7side | GTGACTGGAGTTCAGACGTGTGCTCTTCCGATCT | CCGYCAATTCMTTTRAGTTT | 20 |
| Uni907R2N_TruSecP7side | GTGACTGGAGTTCAGACGTGTGCTCTTCCGATCT | CCGYCTATTCCTTTGAGTTT | 1 |
| Uni907R3N_TruSecP7side | GTGACTGGAGTTCAGACGTGTGCTCTTCCGATCT | CCGYCAATTTCTTTRAGTTT | 1 |
| Uni907R4N_TruSecP7side | GTGACTGGAGTTCAGACGTGTGCTCTTCCGATCT | CCGYCAATTCCCTTRAGTTT | 1 |
| Uni907R5N_TruSecP7side | GTGACTGGAGTTCAGACGTGTGCTCTTCCGATCT | CCGYCAATTCCTTMAAGTTT | 1 |
| Uni907R6N_TruSecP7side | GTGACTGGAGTTCAGACGTGTGCTCTTCCGATCT | CCGCCAATTCCTTTGAATTT | 1 |

F: forward primer; R: reverse primer
